# Antagonistic nanobodies reveal mechanism of GSDMD pore formation and unexpected therapeutic potential

**DOI:** 10.1101/2023.04.20.537718

**Authors:** Lisa D.J. Schiffelers, Sabine Normann, Sophie C. Binder, Elena Hagelauer, Anja Kopp, Assaf Alon, Matthias Geyer, Hidde L. Ploegh, Florian I. Schmidt

## Abstract

Activation of various inflammasomes converges on the cleavage of gasdermin D (GSDMD) by pro-inflammatory caspases, followed by oligomerization of the N-terminal domain (GSDMD^NT^) and the assembly of pores penetrating target membranes. Yet, it remained unclear what triggers the conformational changes that allow membrane insertion, as methods to study pore formation in living cells were limited. We raised nanobodies specific for human GSDMD and found two nanobodies that prevent pyroptosis and IL-1β release when expressed in the cytosol of human macrophages. Nanobody binding to GSDMD^NT^ blocked its oligomerization, while inflammasome assembly and GSDMD processing itself were not affected. The nanobody-stabilized monomers of GSDMD^NT^ partitioned into the plasma membrane, suggesting that pore formation is initiated by insertion of monomers, followed by oligomerization in the target membrane. When GSDMD pore formation was inhibited, cells still underwent caspase-1-dependent apoptosis, likely due to the substantially augmented caspase-1 activity. This hints at a novel layer of regulation of caspase-1 activity by GSDMD pores. Moreover, we revealed the unexpected therapeutic potential of antagonistic GSDMD nanobodies, as recombinant nanobodies added to the medium prevented cell death by pyroptosis, likely by entering through GSDMD pores and curtailing the assembly of additional pores. GSDMD nanobodies may thus be suitable to treat the ever-growing list of diseases caused by activation of the (non-) canonical inflammasomes.

## Introduction

Gasdermin D (GSDMD) is a pore forming protein that perforates the plasma membrane to execute pyroptotic cell death. It is thus considered the key effector protein of the inflammasome pathway^1–4^. Canonical inflammasomes are multiprotein complexes comprised of sensor proteins that oligomerize and recruit the adapter protein ASC as well as pro-inflammatory caspase-1^5^. Distinct inflammasome sensors are activated by different pathogen- or danger-associated molecular patterns (DAMPs). Human NAIP/NLRC4 is activated when NAIP binds to the needle proteins of bacterial type III secretion systems, such as *Shigella flexneri MxiH,* and subsequently initiates oligomerization of NLRC4^6^. NLRP3 is an indirect sensor for potassium efflux and perturbations of intracellular homeostasis^7^. The ensuing activation of caspase-1 is not only responsible for the maturation of pro-inflammatory cytokines IL-1β and IL-18, but also for the cleavage of GSDMD in its interdomain linker. This releases the N-terminus (GSDMD^NT^) from the control of the autoinhibitory C-terminus (GSDMD^CT^), allowing GSDMD^NT^ to assembles pores in the plasma membrane^1–3^. As a result, the plasma membrane becomes permeable to DNA intercalating dyes such as propidium iodide or DRAQ7 and the mature cytokines IL-1β and IL-18 are released^1,3,8^. GSDMD pore formation has recently been found to be enhanced by reactive oxygen species (ROS), which likely mediate oxidative modification of cysteine 192 in murine GsdmdD (corresponding to cysteine 191 in human GSDMD)^9,10^. Eventually, the entire cell ruptures and releases larger cytosolic components into the cellular environment, including tetrameric lactate dehydrogenase (LDH) and pro-inflammatory DAMPs that further promote inflammation. Membrane rupture itself seems to depend on the cell-surface protein Ninjurin-1, which oligomerize in the plasma membrane after GSDMD pore formation^11^.

GSDMD pores reconstituted *in vitro* are composed of 31-34 monomers, forming a pore with an estimated inner diameter of 22 nm^12^. After cleavage, GSDMD^NT^ undergoes drastic conformational changes: two extension domains, composed of short beta sheets and helices, transform into two beta hairpins with extended beta sheets, which constitute the membrane spanning pore upon oligomerization^13,14^. It remained unclear if loss of GSDMD^CT^ is sufficient to trigger these conformational changes, or whether they only occur in concert with oligomerization. Atomic force microscopy suggests that GSDMD^NT^ forms oligomers of different sizes in artificial membranes; smaller slits or arcs were observed to grow into symmetric pores^15^. On the other hand, pore-like structures of human GSDMD and murine GsdmA3 composed of oligomerized globular GSDMD^NT^/GsdmA3^NT^ protomers that do not penetrate the target membrane were observed in samples of purified pores by electron microscopy and on supported lipid membranes by atomic force microscopy^12,13,16^. The authors thus speculated that oligomerization of a prepore precedes conformational changes that lead to membrane penetration^12,13^. Apart from assays reporting the permeability of the plasma membrane to different dyes or cell death, pore formation of endogenous GSDMD had not been studied in molecular detail in living cells, largely due to the lack of suitable tools^8^. Pyroptotic cells are very delicate and are not compatible with staining methods involving fixation and multiple washing steps. Moreover, upon inflammasome activation, fluorescent derivatives of GSDMD^NT^ were barely observed in the plasma membrane, but mostly in intracellular compartments or structures^17^.

To provide more insights into GSDMD pore formation in live cells, we generated nanobodies against the human GSDMD protein. Nanobodies are single domain antibodies derived from the variable domain of heavy chain-only antibodies (VHH) present in *camelids*^18^. Due to their small size, specificity, and functionality in the cytosol, they present themselves as useful tools to study target proteins in living cells^19^. We identified two antagonistic GSDMD nanobodies that inhibit pyroptosis and IL-1β release by blocking oligomerization of GSDMD^NT^. As nanobody-bound GSDMD^NT^ still partitions into the plasma membrane, we concluded that monomeric GSDMD^NT^ exhibits a suitable conformation to insert into the plasma membrane and only oligomerizes after insertion. We discovered an unexpected layer of negative caspase-1 regulation by functional GSDMD pores and found that the inhibitory nanobodies show great potential in preventing inflammatory cell death in primary human macrophages when administered to the extracellular environment. This is of particular interest since GSDMD is linked to an ever-growing list of (auto)inflammatory, metabolic, and neurodegenerative diseases and cancer, and is thus an eminent drug target^20–22^.

## Results

### Identification of GSDMD-specific nanobodies

Specific inhibitors of GSDMD are scarce and apart from complete knockouts and overexpressed point mutants, no tools were available to perturb GSDMD to study pore formation in molecular detail in living cells. To overcome these shortcomings, we raised nanobodies against the human GSDMD protein. An alpaca (*Vicugna pacos*) was immunized with bacterially expressed recombinant full length GSDMD. We employed phage display to positively select GSDMD-specific VHHs, yielding six hits that substantially differ in their complementarity determining regions (CDRs) (Figure 1, A and B). ELISA experiments with decreasing concentrations of the nanobodies confirmed their specificity for GSDMD (Figure 1C). To test binding of the six GSDMD-specific nanobodies to GSDMD in the cytosol of living cells, we performed LUMIER assays in which HEK293T cells were co-transfected with expression vectors for HA-tagged nanobodies (VHH-HA) and for fusions of different variants of GSDMD to Renilla luciferase (GSDMD-Renilla). HA-tagged nanobodies were subsequently immunoprecipitated from lysates. If the nanobody binds to GSDMD in the cells, GSDMD-Renilla is co-immunoprecipitated, resulting in a luminescent signal after addition of the Renilla substrate coelenterazine-h. This confirmed GSDMD binding of VHH_GSDMD-1_, VHH_GSDMD-2_, VHH_GSDMD-3_ and VHH_GSDMD-5_ in the cytosol (Figure 1D). To determine the domain of GSDMD bound by the nanobody, we also included fusions of GSDMD^NT^ and GSDMD^CT^ to Renilla. As overexpression of GSDMD^NT^ alone kills cells by pyroptosis, we used full length and the N-terminal domain of GSDMD mutant 4A, which no longer binds to membranes and does not cause pyroptosis^23^. VHH_GSDMD-1_ and VHH_GSDMD-2_ clearly bind the N-terminal domain of GSDMD, while binding of VHH_GSDMD-3_ and VHH_GSDMD-5_ was affected by the 4A mutation and no clear conclusion was possible. No binding to murine GsdmdD was observed (data not shown).

**Fig. 1.**
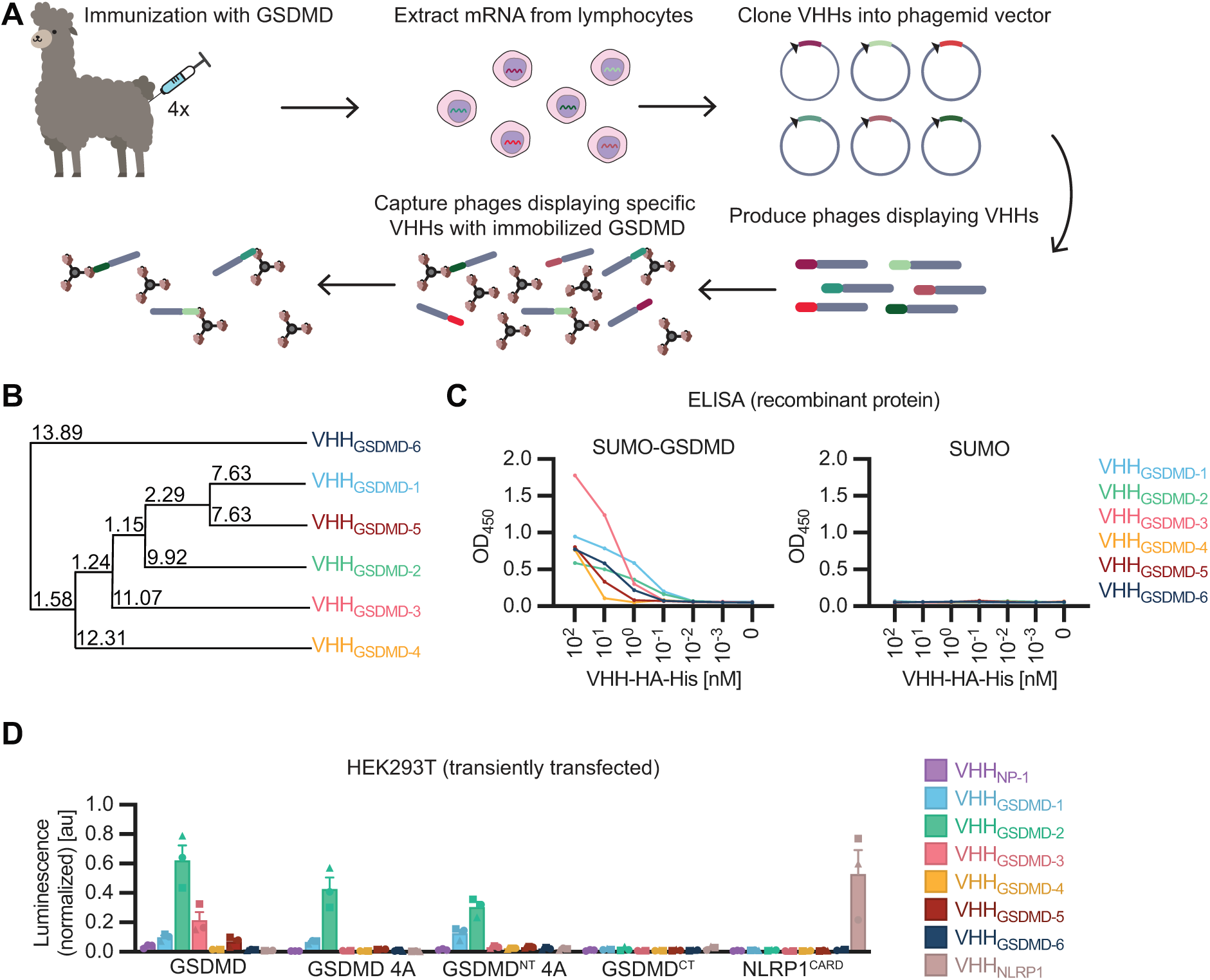
Identification of GSDMD-specific nanobodies. (**A**) Scheme of alpaca immunization and GSDMD nanobody (VHH) selection by phage display. (**B)** Average distance tree calculated based on the percentage identity between the selected nanobody sequences. **(C)** ELISA with recombinant nanobodies: SUMO-GSDMD or control protein SUMO was immobilized on ELISA plates and binding of the indicated concentration of HA-His-tagged nanobodies was quantified by ELISA with anti-HA HRP. **(D)** LUMIER assay: HEK293T cells were co-transfected with expression vectors for the specified HA-tagged nanobodies and the indicated protein-Renilla luciferase fusions. 24 h post transfection, cells were lyzed and VHH-HA was immunoprecipitated with immobilized anti-HA. Coelenterazine-h was added and luminescence of co-purified Renilla luciferase was measured and normalized to luminescence of lysates. Data represent average values (with individual data points) from three independent experiments ± SEM.

### VHH_GSDMD-1_ and VHH_GSDMD-2_ abrogate pyroptosis

Next, we investigated whether the identified nanobodies perturb GSDMD function if expressed intracellularly. To test the effect of the nanobodies on GSDMD^NT^-induced LDH release, HEK293T cells were co-transfected with expression vectors for GSDMD^NT^ and the indicated HA-tagged nanobodies. Interestingly, VHH_GSDMD-1_ and to some extent VHH_GSDMD-2_ inhibited the release of LDH, while the control nanobody and the other GSDMD nanobodies did not affect cell death, suggesting that VHH_GSDMD-1_ and VHH_GSDMD-2_ inhibit pyroptosis (Figure 2A). To validate inhibition of GSDMD pore formation in a more relevant cell type, we generated human myeloid THP-1 cell lines expressing the HA-tagged nanobodies under the control of the strong constitutive EF1α promoter. THP-1 WT cells and cells expressing an unrelated nanobody against the nucleoprotein of influenza A virus (VHH_NP-1_)^24^ were used as negative controls, whereas VHH_ASC_ served as positive control, since it interferes with inflammasome formation and IL-1β release by impairing homotypic interactions between ASC caspase recruitment domains (ASC^CARD^) as well as recruitment of caspase-1^CARD^ by ASC^CARD^ ^25,26^. VHH_GSDMD-1_ and VHH_GSDMD-2_ were expressed at levels similar to the control nanobodies, while VHH_GSDMD-3_ was poorly expressed and excluded from further analyses (Figure S1A). The THP-1 cells were PMA-differentiated into macrophages and activated with either the *Shigella* needle protein MxiH, delivered with the anthrax toxin delivery system to induce NLRC4 inflammasome activation^6^, or with LPS and nigericin to activate the NLRP3 inflammasome^27^. We observed robust LDH and IL-1β release in the control cell lines and confirmed that the responses to both triggers completely depend on caspase-1, as they were abrogated by the caspase-1 inhibitor VX-765 (VX). Responses to LPS and nigericin treatment were mediated by NLRP3, as indicated by the sensitivity to NLRP3 inhibitor CRID3 (Figure 2, B, C, D, and E). Strikingly, both VHH_GSDMD-1_ and VHH_GSDMD-2_ completely shut down the release of LDH (Figure 2, B and C) and IL-1β (Figure 2, D and E) to background levels after NLRC4 as well as NLRP3 inflammasome activation. We next activated inflammasomes and microscopically followed and quantified the uptake of DRAQ7, a membrane-impermeable far-red fluorescent DNA dye, to monitor permeability of the plasma membrane over time. While robust DRAQ7 uptake was detected in WT cells and cells expressing VHH_NP-1_ within 1 h after treatment, no DRAQ7 uptake was apparent in the THP-1 macrophages expressing VHH_GSDMD-1_ or VHH_GSDMD-2_ (Figure 2, F and G). In line with these findings, cells expressing these antagonistic nanobodies do not show any features of pyroptotic cell death, as opposed to the negative controls (Figure 2F). THP-1 macrophages expressing VHH_ASC_ did not allow any DRAQ7 influx after NLRP3 stimulation. This is consistent with the critical role for ASC in NLRP3 inflammasome assembly and caspase-1 recruitment. In contrast, VHH_ASC_ expression did not fully inhibit DRAQ7 uptake after NLRC4 activation, perhaps because ASC inhibition was incomplete or because NLRC4 can directly recruit and activate caspase-1^28^. We conclude that cytosolic expression of VHH_GSDMD-1_ and VHH_GSDMD-2_ completely abrogates GSDMD-mediated effector functions. As GSDMD pore formation after NLRP3 and NLRC4 activation was similarly affected by antagonistic GSDMD nanobodies, subsequent experiments focused on NLRC4 activation.

**Fig. 2.**
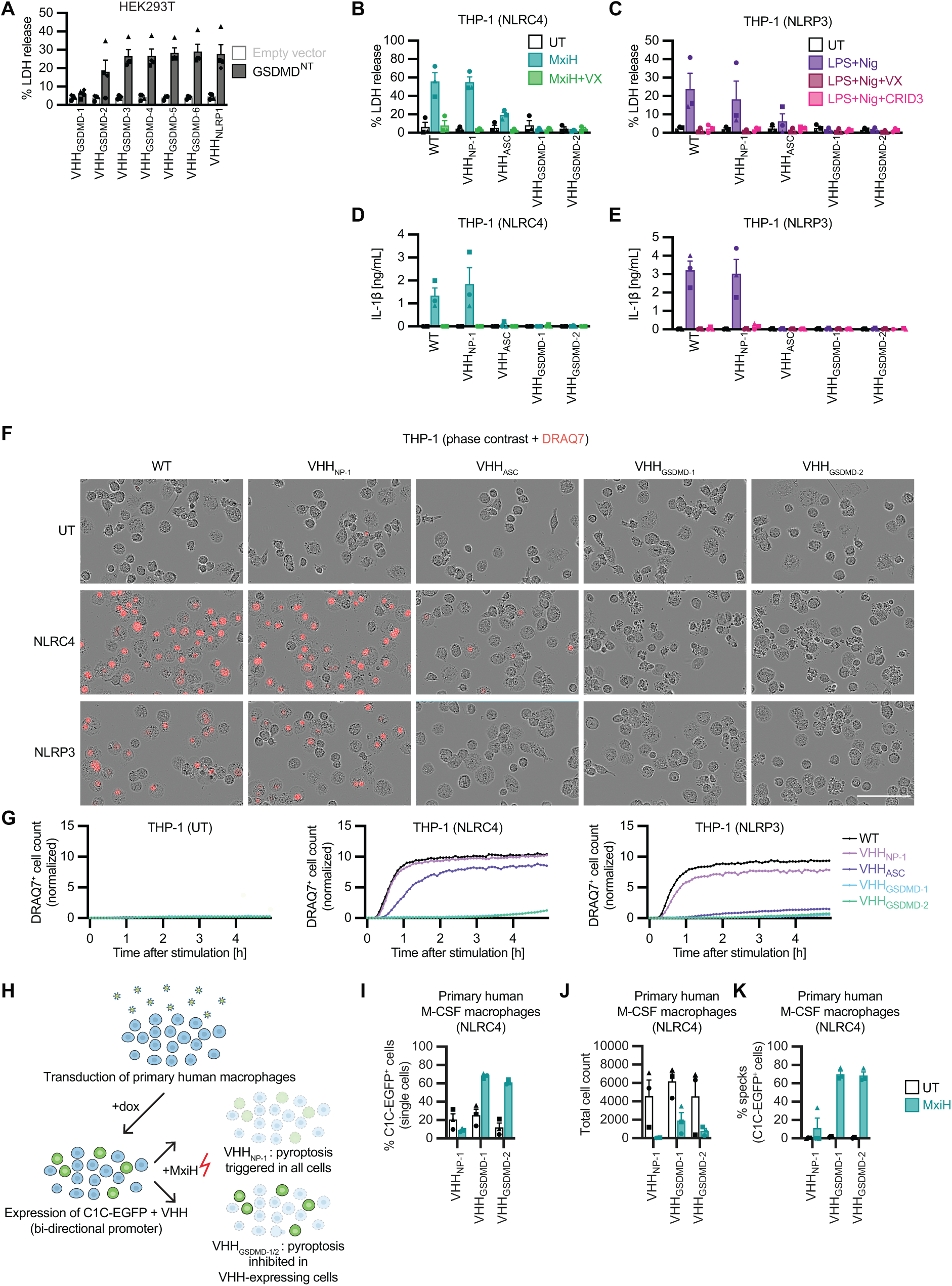
VHH_GSDMD-1_ and VHH_GSDMD-2_ abrogate pyroptosis. (**A**) HEK293T cells were co-transfected with expression vectors for the indicated HA-tagged nanobodies as well as GSDMD^NT^ or empty vector. LDH release was measured 24 h post transfection and normalized to cells lysed in 1% Triton X-100 (n=4). **(B-E)** PMA-differentiated THP-1 macrophages constitutively expressing the indicated HA-tagged nanobodies or WT controls were stimulated with 1.0 µg/mL PA and 0.1 µg/mL LFn-MxiH (MxiH) for 1 h to activate NLRC4 (B, D), or with 200 ng/mL ultrapure LPS for 3 h and 10 µM nigericin (Nig) for 1 h to activate NLRP3 (C, E), in the presence of 40 µM VX-765 (VX) or 2.5 µM CRID3 where indicated. LDH release was measured as in A (B, C), and IL-1β in the supernatant was measured by Homogeneous Time Resolved Fluorescence (HTRF) (D, E). **(F, G)** PMA-differentiated THP-1 macrophages were stimulated with NLRC4 and NLRP3 activators as described above, but in the presence of 100 nM DRAQ7. DRAQ7 uptake was monitored over 5 h in an Incucyte Live-Cell Imaging system. Representative images (F) and graphs (G) (of n=3) after 1 h of normalized DRAQ7 uptake are displayed. Scale bar: 100 µm. **(H)** Overview of transduction of primary human macrophages with lentivirus particles packaging Vpx-Vpx and encoding C1C-EGFP and the different nanobodies under the control of a bi-directional doxycycline (dox)-inducible promoter. Stimulation with NLRC4 activator MxiH triggers cell death by pyroptosis, unless the expressed nanobodies inhibit GSDMD pore formation, which leads to the enrichment of the respective transduced (C1C-EGFP-positive) cells. **(I-K)** Primary M-CSF-differentiated monocyte-derived human macrophage were transduced with lentivirus particles encoding C1C-EGFP and the indicated nanobody. 24 h post transduction, gene expression was induced with dox and 24 h later, cells were treated with NLRC4 activator MxiH as in B and D. 1 h post treatment, cells were harvested, fixed, and analyzed by flow cytometry to determine the fraction of C1C-EGFP^+^ and thus VHH-expressing cells (I), total cell count over 30 s (J), and the fraction of C1C-EGFP^+^ cells assembling ASC specks (K). Data represent average values (with individual data points) from three independent experiments or donors ± SEM, unless mentioned otherwise.

We next investigated the functionality of the inhibitory GSDMD nanobodies in primary human M-CSF- and GM-CSF-differentiated macrophages. For this purpose, primary human macrophages were transduced with lentivirus encoding the nanobody of interest in addition to our previously described fluorescent inflammasome reporter caspase-1^CARD^-EGFP (C1C-EGFP), which is recruited to ASC specks via the CARD of caspase-1^29^ (Figure 2H). To overcome restriction by SAMHD1 in macrophages, lentivirus was produced in cells expressing a fusion protein of SIVmac251 Vpx and HIV-1 NL4.3 Vpr. Vpx-Vpr is packaged into lentivirus particles as Vpr binds to the structural protein Gag, and thus delivers Vpx into target cells, which mediates the Cullin-4a-mediated proteasomal degradation of SAMHD1^30,31^. This led to a transduction efficiency of 10-40%, as measured by flow cytometry (Figure 2I, Figure S1B). Upon treatment with the NLRC4-activating trigger MxiH, we observed a strong reduction in cell counts, as pyroptotic cells are too fragile to survive processing for flow cytometry (Figure 2J, Figure S1C). Loss of cells was almost complete in cells transduced with lentivirus expressing control nanobodies and less prominent in cells transduced with lentivirus expressing antagonistic GSDMD nanobodies. At the same time, we observed that macrophages expressing antagonistic VHH_GSDMD-1_ or VHH_GSDMD-2_ and C1C-EGFP preferentially survived, as EGFP-positive cells now comprise 60-70% of the single cell population (Figure 2I, Figure S1B), whereas cells expressing control nanobody VHH_NP-1_ did not provide a survival benefit. This indicates that only the cells expressing the antagonistic VHHs could survive the lethal trigger, and hence VHH_GSDMD-1_ and VHH_GSDMD-2_ prevent pyroptosis in primary human macrophages. To verify that MxiH treatment induced inflammasome assembly, we quantified recruitment of C1C-EGFP to ASC specks by flow cytometry as described before^29,32^. Briefly, characteristic changes in the width and height of the C1C-EGFP signal during speck formation allow detection of cells with an ASC speck as a separate population. Robust inflammasome activation was observed in the majority of MxiH-treated cells that survived NLRC4 activation due to the expression of antagonistic GSDMD nanobodies. This confirms that the surviving cells did respond to MxiH and survival was not due to any impairment in inflammasome assembly (Figure 2K, Figure S1D).

Altogether, these results show a potent pyroptosis-inhibiting effect of VHH_GSDMD-1_ and VHH_GSDMD-2_ in both THP-1 macrophages and primary human macrophages upon NLRP3 and NLRC4 inflammasome activation.

### Antagonistic nanobodies prevent the oligomerization of GSDMD^NT^, but still allow membrane localization

We were wondering what the mechanism of pyroptosis inhibition by the GSDMD nanobodies could be. To rule out any effect on the assembly of inflammasomes, we analyzed the formation of ASC specks in THP-1 macrophage cell lines, which -in addition to the different VHH-EGFP fusions- also inducibly expressed the C1C-mCherry inflammasome reporter. Flow cytometry analysis upon MxiH delivery or stimulation with LPS and nigericin (in presence of VX to prevent pyroptosis and therefore cell loss) proved that the nanobodies do not influence formation of ASC specks (Figure 3A, Figure S2A).

**Fig. 3.**
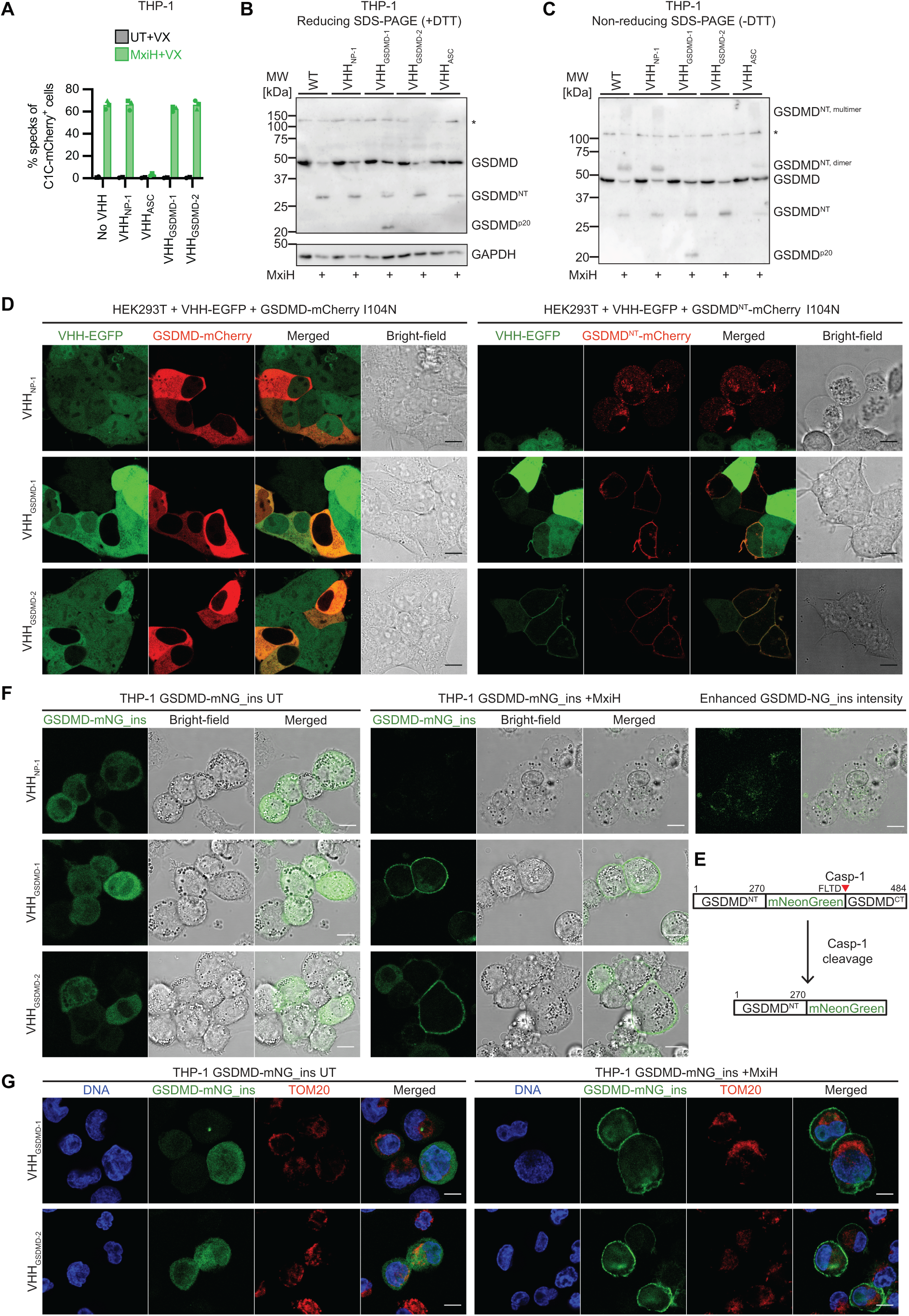
Antagonistic nanobodies prevent the oligomerization of GSDMD^NT^, but still allow membrane localization. (**A**) THP-1 cell lines expressing C1C-mCherry (dox-inducible) as well as the indicated VHH-EGFP fusions (constitutively) were differentiated with PMA, treated with dox for 24 h, and subjected to stimulation with NLRC4 agonist MxiH for 1 h as described in Fig. 2B, in presence of 40 µM VX. Cells were harvested and ASC specks were quantified by flow cytometry. Data represent average values (with individual data points) from three independent experiments ± SEM. **(B, C)** PMA-differentiated THP-1 macrophages expressing the indicated HA-tagged nanobodies were stimulated with MxiH for 1 h as described in Fig. 2B. Cells were lysed in SDS-PAGE buffer with 100 mM DTT (B) or no reducing agent (C), and subjected to SDS-PAGE and immunoblot with GSDMD and GAPDH antibodies. Representative immunoblot of at least three independent experiments are displayed. **(D)** HEK293T cell stably expressing the indicated VHH-EGFP fusions were transfected with expression vectors for GSDMD-mCherry I104N (left) or GSDMD^NT^-mCherry I104N (right) and analyzed by live-cell confocal imaging. Data representative of three independent experiments are shown. Scale bar = 10 µm. **(E)** Schematic representation of GSDMD-mNeonGreen_ins (GSDMD-mNG_ins) before and after cleavage by caspase-1. **(F, G)** PMA-differentiated THP-1 cells constitutively expressing GSDMD-mNG_ins and the indicated HA-tagged nanobodies were stimulated with MxiH for 30 minutes (G) or 1 h (F) as described in Fig. 2B. Living (F) or fixed (G) cells were recorded by confocal microscopy; DNA and mitochondria in fixed cells were stained using Hoechst and TOM20 antibodies with fluorescent secondary antibodies. Representative images are displayed. Scale bar = 10 µm.

Next, we analyzed whether binding of VHH_GSDMD-1_ or VHH_GSDMD-2_ altered GSDMD levels or GSDMD processing by caspase-1. Lysates of MxiH-treated THP-1 macrophages were separated by SDS-PAGE under reducing conditions and analyzed by immunoblot. No differences in GSDMD expression and cleavage into GSDMD^NT^ were detected (Figure 3B). It had previously been shown that GSDMD oligomers appeared as high-molecular weight bands by SDS-PAGE under non-reducing conditions^23,33^. We thus analyzed THP-1 cell lysates separated by SDS-PAGE in the absence of DTT. In MxiH-treated WT THP-1 cells and cells expressing control nanobodies, we observed the formation of dimers and higher order oligomers in absence of DTT, which were no longer present in the lysates from VHH_GSDMD-1_ or VHH_GSDMD-2_ -expressing cells (Figure 3C). This demonstrates that the nanobodies interfere with the oligomerization of GSDMD^NT^.

Having established a system in which antagonistic nanobodies stabilize monomeric GSDMD^NT^ by preventing oligomerization, we were curious if the monomeric protein would be sufficient to insert into membranes, or whether this step required oligomerization. We therefore transfected HEK293T cells stably expressing VHH-EGFP fusions with expression vectors for full length GSDMD-mCherry or GSDMD^NT^-mCherry with the attenuating mutation I104N^2,4,34^ and followed the localization by live cell confocal microscopy. The GSDMD-mCherry signal was evenly distributed in the cytosol irrespective of the nanobody expressed, confirming that full length GSDMD did not insert into the plasma membrane as expected (Figure 3D). When GSDMD^NT^-mCherry I104N was expressed in HEK293T cells expressing control nanobody VHH_NP-1_, we observed that most cells appeared pyroptotic, while fluorescence was mostly found in internal structures. This suggests that pores of GSDMD^NT^ do not predominantly accumulate in the plasma membrane. It is possible that initial pores in the plasma membrane are rapidly removed, e.g., by membrane repair processes involving shedding of membrane vesicles^35^. It cannot be ruled out that some GSDMD^NT^ is also recruited to other cellular membranes, as proposed earlier, mostly based on indirect evidence^36–39^. In contrast, when GSDMD^NT^ was co-expressed in cells with VHH_GSDMD-1_ or VHH_GSDMD-2_, GSDMD^NT^-mCherry almost completely partitioned into the plasma membrane, where it co-localized with VHH-EGFP (Figure 3D). The cells no longer showed signs of pyroptosis, in line with the expected inhibition of GSDMD pore formation. Monomeric nanobody-bound GSDMD^NT^ can thus insert into the plasma membrane, indicating that the required conformational changes for membrane integration occur in monomeric GSDMD after removal of the auto-inhibitory C-terminus. This implies that GSDMD^NT^ only oligomerizes after inserting into the plasma membrane, and therefore most likely does not need to go through a prepore state comprised of oligomeric GSDMD^NT^ that is not inserted into the membrane.

We had unsuccessfully tried to reproducibly visualize GSDMD pores in pyroptotic cells after inflammasome activation in THP-1 cells inducibly expressing GSDMD with mNeonGreen inserted after amino acid 270 (GSDMD-mNeonGreen_ins) (Figure 3E)^17^. Insertion of the fluorescent protein between the GSDMD^NT^ and the caspase-1 cleavage site yields GSDMD^NT^-mNeonGreen after caspase-1 cleavage. Similar to our observations in HEK293T cells expressing GSDMD^NT^-mCherry, most of GSDMD^NT^-mNeonGreen generated by caspase-1 cleavage localized to internal structures, but not to the plasma membranes. Given that VHH_GSDMD-1_ and VHH_GSDMD-2_ stabilized GSDMD^NT^ monomers in the plasma membrane in HEK293T cells, we wondered if the antagonistic nanobodies also stabilized GSDMD^NT^-mNeonGreen cleaved after inflammasome-mediated caspase-1 activation in relevant cell types. We therefore generated THP-1 cells expressing VHH_GSDMD-1_-HA or VHH_GSDMD-2_-HA in addition to GSDMD-mNeonGreen_ins. When inflammasome activation was triggered in those cell lines, cells did not exhibit morphological features of pyroptosis and GSDMD^NT^-mNeonGreen localized to the plasma membrane (Figure 3F, Figure S2B). This confirmed that GSDMD^NT^ released by cleavage of full length GSDMD also inserted into the plasma membrane as a monomer before pores were formed by oligomerization. Importantly, GSDMD^NT^-mNeonGreen was not clearly detected in specific internal structures and no co-localization with mitochondria was apparent (Figure 3G).

Monitoring GSDMD^NT^ by fluorescence microscopy therefore allows two additional relevant conclusions: 1) Stabilized GSDMD^NT^ monomers predominantly localize to the plasma membrane, rendering it less likely that insertion into other organelles such as mitochondria contributes to the initial phases of pyroptosis. 2) Fluorescent GSDMD^NT^ is only observed in internal vesicular structures when GSDMD^NT^ is able to form pores, i.e., in cells with control nanobodies, suggesting that the observed structures result from the internalization of GSDMD pores and thus endocytic repair mechanism that remove GSDMD pores.

In conclusion, our inhibitory GSDMD nanobodies interfere with GSDMD^NT^ oligomerization, while cleavage and membrane localization in living cells are not affected. We propose that monomeric GSDMD^NT^ is able to directly insert into the plasma membrane and build up the pore monomer by monomer within target membranes. This implies that there is no need to oligomerize prepores before membrane insertion.

### Inhibition of pore formation by antagonistic GSDMD nanobodies augments caspase-1 activity and triggers caspase-1-dependent apoptosis

When analyzing THP-1 macrophages expressing different VHH-EGFP fusions in combination with the C1C-mCherry inflammasome reporter with live cell microscopy, we observed inflammasome assembly in presence of VHH_GSDMD-1_ and VHH_GSDMD-2_ as indicated by ASC speck formation within 1 h post treatment (Figure 4A). Interestingly, cells with ASC specks exhibited blebs or were fragmented into multiple membrane-surrounded structures – morphologies more typically associated with apoptosis and apoptotic bodies (also see Figure 2F). Cells with ASC specks expressing the VHH_NP-1_ control, however, were round-up with a ‘balloon-like’ morphology as expected for cells undergoing pyroptosis (Figure 4A, Figure 2F). Similar apoptotic morphologies could be observed for the primary human macrophages transduced with antagonistic GSDMD nanobodies and C1C-EGFP as described above (Figure S3A, Figure 2H).

**Fig. 4.**
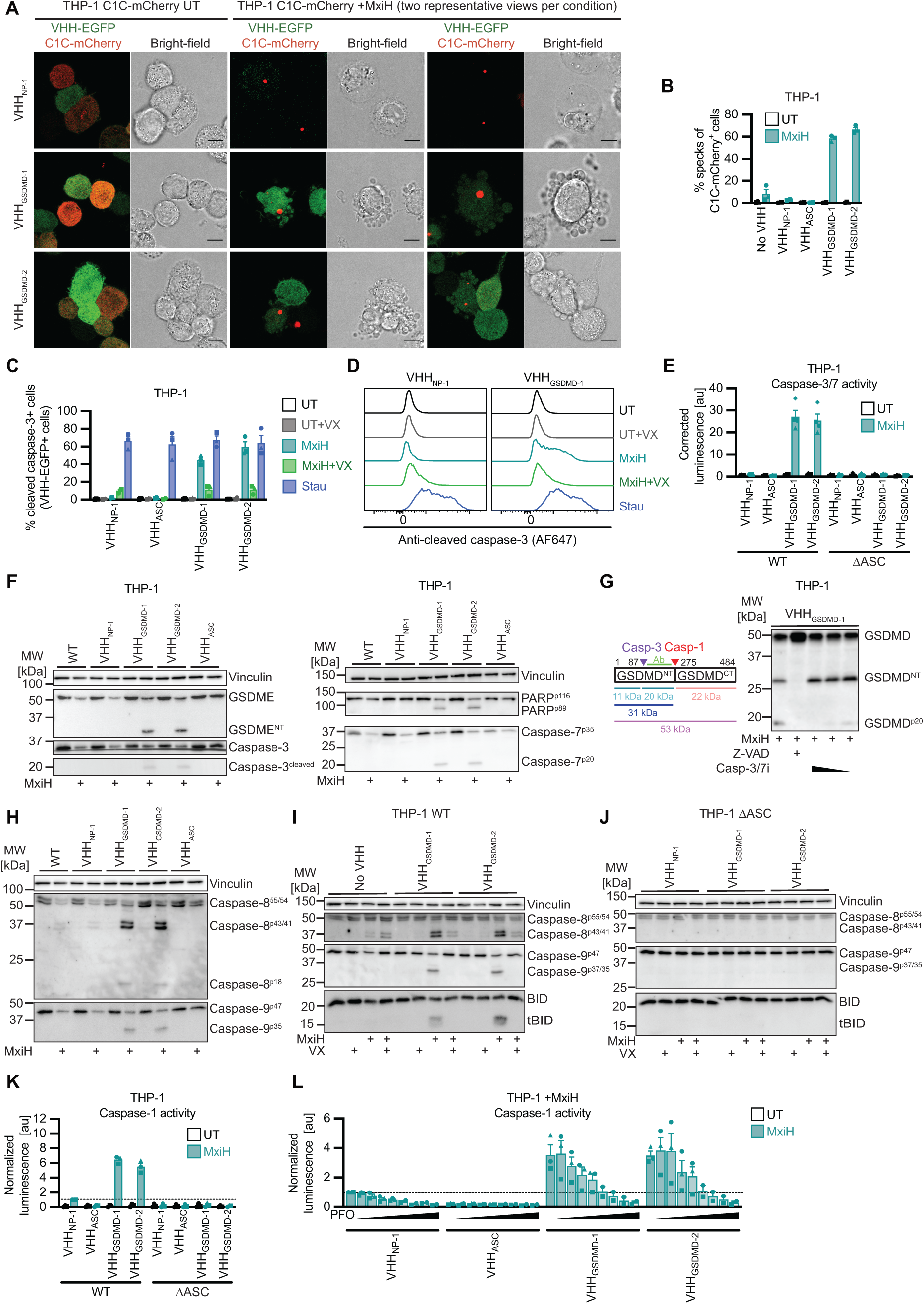
Inhibition of pore formation by antagonistic GSDMD nanobodies augments caspase-1 activity and triggers caspase-1-dependent apoptosis. **(A-D)** THP-1 cell lines expressing C1C-mCherry (dox-inducible) as well as the indicated VHH-EGFP fusions (constitutively) were differentiated with PMA, treated with dox for 24 h, and stimulated with NLRC4 agonist MxiH for 1 h as described in Fig. 2B, or with 5 µM staurosporine (Stau) for 20 h to trigger apoptosis. Stimulation was performed in the absence or presence of VX as indicated. (A) Cells were recorded by live cell confocal microscopy and images representative of three independent experiments are displayed (two fields of view for treated samples). Scale bar = 10 µm. (B) Cells were harvested and analyzed by flow cytometry to quantify C1C-mCherry specks. (C, D) Cells were harvested, stained with antibodies specific for cleaved caspase-3 and Alexa Fluor 647 (AF647)-coupled secondary antibodies, and the fraction of cells positive for cleaved caspase-3 was quantified by flow cytometry (C). Representative histograms of two exemplary cell lines with the indicated treatments are presented in (D), the remaining histograms are displayed in Fig. S4C. (E-L) PMA-differentiated THP-1 WT (E-I, K, L) or THP-1 ΔASC (E, J, K) cells constitutively expressing the indicated HA-tagged nanobodies were stimulated with MxiH for 1 h as described in Fig. 2B. Experiments were performed in the presence of 40 µM VX, 50 µM Z-VAD, caspase-3/7 inhibitor (30, 20, and 4 µM), or PFO (0, 5, 10, 20, 30, 60, 120, 240, and 480 ng/mL), as indicated (G, I, J, L). Cells and supernatants were harvested to measure caspase-3/7 (E) or caspase-1 (K, L) activity using Caspase-Glo assays. Luminescence was corrected for total cell numbers per sample using CTB values (E, K, L); caspase-1 activity was normalized to MxiH-treated cells expressing control VHH_NP-1_ (indicated as dashed line) (K, L). Caspase-1 activity of untreated samples from the same experiments as (L) is displayed in Fig. S4I. Data represent average values (with individual data points) from three (E, K, L) independent experiments ± SEM. (F-J) Cell lysates were separated by SDS-PAGE and analyzed by immunoblot with the indicated antibodies. Immunoblots with caspase-3 and cleaved caspase-3 antibodies were prepared from the same (cut) membrane and developed separately, with a longer exposure for caspase-3^cleaved^ (F). Caspase-3 (Casp-3) and caspase-1 (Casp-1) cleavage sites in GSDMD, resulting fragments, and the peptide used to raise the GSDMD antibody (Ab) are indicated next to the GSDMD immunoblots (G). Data represent average values (with individual data points) from three independent experiments ± SEM for all flow cytometry and caspase activity assays. Representative immunoblots or microscopy images of at least three independent experiments are displayed.

When comparable samples after NLRC4 or NLRP3 activation were analyzed by flow cytometry, C1C-EGFP specks were only detected in cells expressing VHH_GSDMD-1_ and VHH_GSDMD-2_, but not in cells expressing control nanobodies or no nanobodies (Figure 4B, Figure S3B). This confirms that pyroptotic cells were ruptured during sample processing as observed before, while apoptotic cells could be analyzed by flow cytometry. Of note, we had treated cells with caspase-1 inhibitor VX to quantify inflammasome assembly in all cells irrespective of pyroptosis in earlier experiments (Figure 3A).

To probe for *bona fide* apoptosis, we next stained the different PMA-differentiated THP-1 cell lines for cleaved caspase-3 after MxiH treatment for 1 h and quantified the fraction of cells positive for cleaved caspase-3 by flow cytometry. Staurosporine treatment for 20 h was used as a positive control and resulted in more than 60% of the cells positive for cleaved caspase-3 (Figure 4C). Both VHH_GSDMD-1_ and VHH_GSDMD-2_ expressing cells treated with MxiH showed a clear population of cells positive for cleaved caspase-3, indicating that the apoptotic effector caspase-3 is active (Figure 4, C and D, Figure S3C). Interestingly, caspase-3 activation seems to be caspase-1-dependent since it was strongly reduced in presence of VX. Direct activation of caspase-8 by recruitment and autoproteolytic activation on ASC specks had been reported earlier^40–42^. As caspase-3 activation was largely blocked by VX, caspase-1-independent activation of caspase-8 does not seem to have a major contribution to caspase-3 activation. Yet the residual fraction of cells positive for caspase-3 cleavage after VX treatment may result from direct activation of caspase-8 on ASC specks. In line with this interpretation, residual caspase-3 activation was no longer observed when cells expressed VHH_ASC_, which prevents the formation of ASC specks^25^. Flow cytometry analysis of caspase-3 cleavage in THP-1 ΔASC cells expressing the different VHHs also showed complete loss of caspase-3 activity, indicating dependence on ASC speck formation (Figure S3, D and E). While NLRC4^CARD^ had been reported to directly recruit caspase-1^CARD^ in the absence of ASC^28^, no LDH release was observed in THP-1 ΔASC cells treated with MxiH, indicating that ASC-independent caspase-1 activation did not contribute in our experimental conditions (Figure S3F).

Flow cytometry does not allow the measurement of pyroptotic cells and flow-cytometry-based analysis may thus fail to detect cleaved caspase-3 in the control cell lines that still undergo pyroptosis. To measure caspase-3 activity independent of cell death or rupture, we performed caspase Glo assays to measure the activity of caspase-3/7 in THP-1 macrophages upon MxiH treatment. Here, caspase activity is determined in lysates derived from the cells and the supernatant, whereby the caspase-3-specific peptide DEVD is cleaved to render a substrate available to luciferase. Strong caspase-3/7 activity was observed in MxiH-treated cells expressing VHH_GSDMD-1_ or VHH_GSDMD-2_, but not in cells expressing control nanobodies. Again, this activity was completely dependent on ASC, as no caspase-3/7 activity was observed in ASC knockout cells (Figure 4E).

Analysis of THP-1 macrophage cell lysates by immunoblot confirmed cleavage of caspase-3, caspase-7, and the caspase-3 substrates PARP and GSDME specifically in those samples with caspase-3 activity, i.e., in cells in which antagonistic GSDMD nanobodies prevented pore formation (Figure 4F). Remarkably, cleavage of GSDME in these cells does not seem to be sufficient to assemble functional GSDME pores, as we did not observe pyroptosis, IL-1β release, or DRAQ7 uptake (Figure 2, C, E, and F). GSDME therefore does not seem to play a major role in the death of VHH_GSDMD_-expressing cells. The GSDMD fragment GSDMD^p20^ in cells expressing VHH_GSDMD-1_ (Figure 3B, Figure 4G) also coincides with enhanced caspase-3 activation and disappears upon addition of a caspase-3/7 inhibitor, suggesting that it represents GSDMD^NT^ cleaved by caspase-3 (Figure 4G). Of note, GSDMD^p20^ is only observed in cell lines expressing VHH_GSDMD-1_ but not VHH_GSDMD-2_, perhaps because access of caspase-3 is occluded by VHH_GSDMD-2_.

Lysates of THP-1 macrophages expressing VHH_GSDMD-1_ and VHH_GSDMD-2_ not only contained cleaved caspase-3, but also processed caspase-8, processed caspase-9, and cleaved tBID (Figure 4H, Figure S3, G and H), indicating the activation of both the intrinsic and extrinsic apoptosis pathway, or the activation of feedback mechanisms involving the upstream caspases. Caspase-1, caspase-8, and caspase-9 can all catalyze the cleavage of caspase-3. As we found that caspase-3 activity depends on caspase-1 and ASC, caspase-1 activated at the inflammasome seems to be the key regulator of the alternative cell death program. Coherently, we found that caspase-8 and caspase-9 activity as well as tBID cleavage are largely dependent on caspase-1 activity and inflammasome formation, as their processing is strongly reduced in presence of VX and in THP-1 ΔASC macrophages (Figure 4, I and J, Figure S3G). Only for caspase-8 there is some residual processing that can also be seen in the absence of GSDMD VHHs (Figure 4I). This caspase-8 activation is completely ASC dependent since it is absent in the THP-1 ΔASC cells (Figure 4J), suggesting that a small portion of the caspase-8 is cleaved at the ASC speck, independent of caspase-1 as concluded above^40–42^.

We next quantified the caspase-1 activity of THP-1 macrophages upon MxiH stimulation using caspase-1 Glo assays. Surprisingly, we found that caspase-1 activity was increased up to 6-fold in presence of VHH_GSDMD-1_ or VHH_GSDMD-2_ compared to the pyroptotic cells expressing VHH_NP-1_ (Figure 4K). This is remarkable, as the assembly of ASC specks was comparable in all samples (Figure 3A). We therefore hypothesize that the ability to form GSDMD pores has a profound impact on caspase-1 activity, suggesting GSDMD pores downregulate caspase-1 activity in a so far elusive mechanism. Caspase-1 ultimately serves as the master regulator for downstream cell death, as only the enhanced caspase-1 activity observed in the absence of GSDMD pores was sufficient to activate caspase-3 and apoptosis.

The relatively low caspase-1 activity in cells undergoing pyroptosis may be a consequence of ion fluxes and/or the release of caspase-1 into the supernatant through GSDMD pores, even though caspase-1 activity was measured in samples derived from the cells and the supernatant. The altered environment may, e.g., compromise the stability of short-lived active caspase-1. To test the impact of pores on caspase-1 activity, we treated THP-1 macrophages expressing VHH_GSDMD-1_ or VHH_GSDMD-2_ with MxiH in the presence of the pore-forming toxin perfringolysin O (PFO) from *Clostridium perfringens*. PFO forms pores with a diameter of 25-30 nm, i.e., a similar if not slightly larger diameter than GSDMD pores^43^. To avoid additional activation of NLRP3 by potassium efflux through PFO pores, NLRC4 was activated in the presence of NLRP3 inhibitor CRID3. PFO-induced pore formation was confirmed by influx of DRAQ7 (Figure S3J). We indeed observed a dose-dependent reduction in caspase-1 activity upon higher levels of PFO administration, indicating that the mere formation of pores in the cellular membrane reduces the activity of caspase-1 (Figure 4L, Figure S3I).

In summary, we propose that the observed apoptosis in the absence of functional GSDMD pores is completely dependent on inflammasome assembly. The augmented caspase-1 activity observed in the absence of GSDMD pores seems to be central to process apoptotic initiator and effector caspases.

### Recombinant antagonistic GSDMD nanobodies inhibit pyroptosis when administered extracellularly

GSDMD is a highly sought-after drug target, although the development of specific GSDMD inhibitors was not successful to date^44–47^. Therapeutic application of the potent pyroptosis-inhibiting nanobodies described in this study would require efficient delivery of nanobodies into the cytosol of target cells. While nanobodies are unable to cross intact cell membranes, we had previously observed that fluorescently labeled nanobodies can enter pyroptotic cells, likely via passage through GSDMD pores^26^. We therefore speculated that early GSDMD pores may allow the influx of antagonistic GSDMD nanobodies administered from outside. To test this, we recombinantly produced the identified GSDMD nanobodies in bacteria and added increasing concentrations of the purified nanobodies to the culture medium of THP-1 macrophages treated with MxiH as an NLRC4 inflammasome activator. We observed a dose-dependent reduction in LDH release and importantly, higher concentrations of VHH_GSDMD-1_ and VHH_GSDMD-2_ reduced LDH release to background levels. In contrast, addition of increasing amounts of VHH_NP-1_ did not affect the release of LDH (Figure 5A). The secretion of IL-1β was also substantially reduced, although the highest concentrations did not completely abrogate cytokine release (Figure 5B). In accordance with the data provided in *Figure 4*, extracellular administration of the antagonistic GSDMD nanobodies, but not VHH_NP-1_, changed the inflammasome-mediated type of cell death from pro-inflammatory pyroptosis to non-inflammatory apoptosis (Figure 5C). To validate these findings in a physiologically relevant *in vitro* model, we repeated the same experiments in primary human M-CSF macrophages. Here, LDH release was similarly inhibited in a dose-dependent manner; IL-1β release was completely abrogated at the highest concentrations (Figure 5, D and E).

**Fig. 5.**
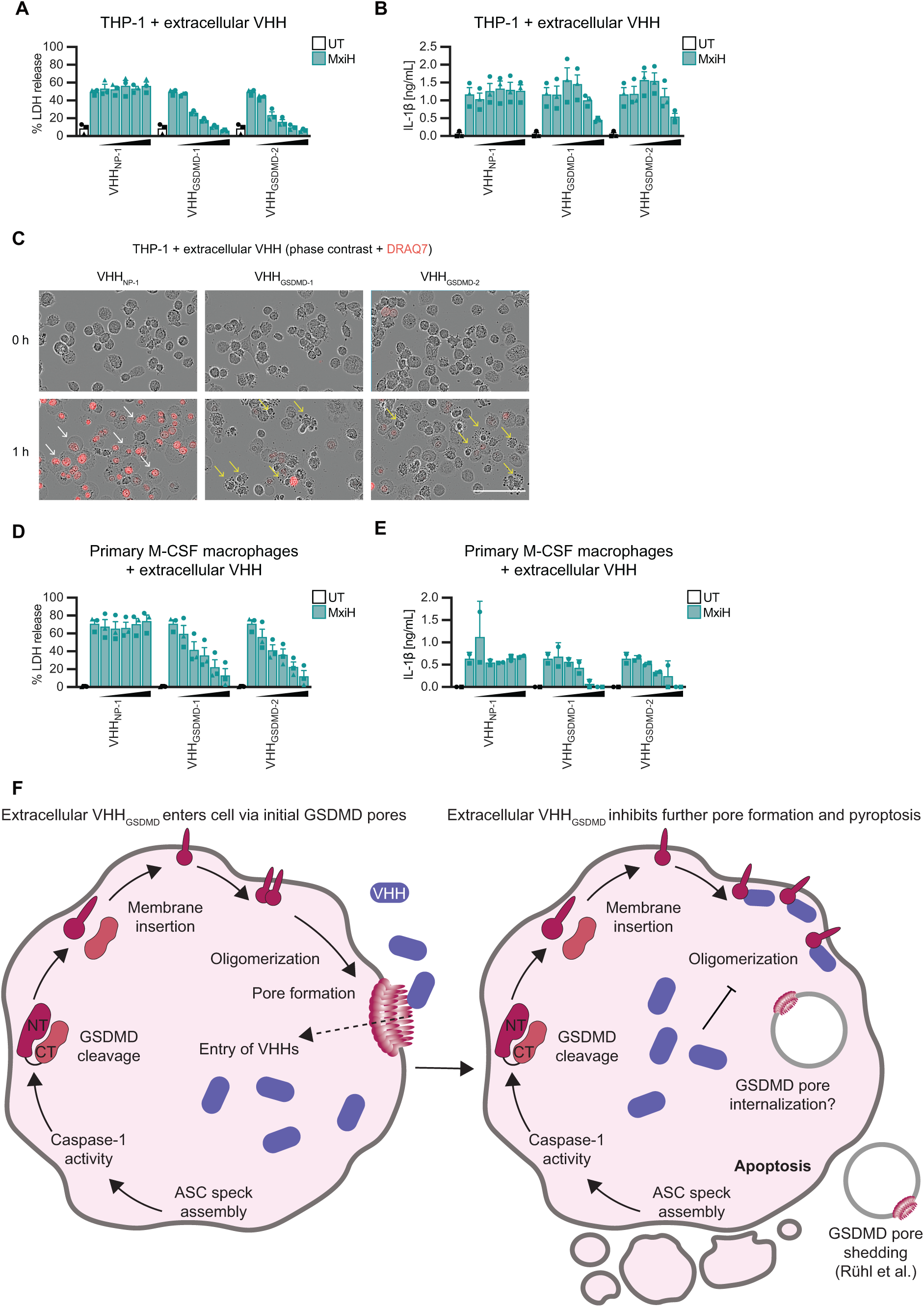
Recombinant antagonistic GSDMD nanobodies inhibit pyroptosis when administered extracellularly. (**A-C**) PMA-differentiated THP-1 cells were treated with MxiH for 1 h as described in Fig. 2B in the presence of increasing concentrations (2, 20, 50, 100, and 200 µg/mL) of the indicated recombinant nanobodies. (A) LDH release was measured and normalized to cells lysed in Triton X-100. (B) IL-1β in the supernatant was quantified by HTRF. (C) THP-1 cells treated in the presence of DRAQ7 were recorded with an Incucyte Live-Cell Imaging system. Representative images are displayed. White arrows indicate pyroptotic cells, yellow arrows indicate apoptotic cells. Scale bar: 100 µm. **(D,E)** M-CSF-differentiated primary human macrophages from independent donors were treated as in Fig. 5A-C, and LDH release (three donors) (D) and IL-1β secretion (two donors) (E) were quantified as before. Data on LDH and IL-1β release represent average values (with individual data points) from three independent experiments or donors unless indicated otherwise ± SEM. **(F)** Model for the inhibition of pyroptosis by antagonistic GSDMD nanobodies added to the extracellular space.

We hypothesize that nanobodies enter cells with inflammasomes upon formation of the first GSDMD pores, before the lytic stage of pyroptosis (Figure 5F). Cytosolic nanobodies may thus prevent any further GSDMD pore assembly, which seems to be sufficient to prevent cell lysis. Based on our experiments with fluorescent GSDMD^NT^ fusions, it is likely that the early GSDMD pores are rapidly removed by membrane repair processes. Initial pore formation may well explain the remaining IL-1β secretion, since the cytokine may still be released through early sublytic GSDMD pores. Altogether, these results show that the nanobodies are potent inhibitors of inflammasome-induced pyroptosis when administered extracellularly, which reveals their interesting therapeutic potential. Importantly, early GSDMD pore formation does not seem to be a terminal event, as cells could still be rescued from pro-inflammatory pyroptosis by antagonistic GSDMD nanobodies.

## Discussion

GSDMD pore formation is the effector mechanism that mediates cell death by pyroptosis as well as the non-conventional secretion of mature IL-1β and IL-18. The function of GSDMD was revealed in seminal loss-of-function screens^1,2^ and the structures of soluble full length GSDMD and GSDMD^NT^ pores were elucidated by X-ray crystallography and electron microscopy^12,14^. Despite these advances, critical molecular aspects of pore formation remained unknown as the process cannot be easily studied in relevant cells, primarily because pyroptotic cells do not weather sample preparation for microscopy and flow cytometry. In this study, we discovered two GSDMD targeting nanobodies, VHH_GSDMD-1_ and VHH_GSDMD-2_, which potently inhibit pyroptosis by preventing the oligomerization of GSDMD^NT^ and thus stabilizing monomeric GSDMD^NT^. Importantly, monomeric GSDMD^NT^ was still able to partition into the plasma membrane. This revealed that cleavage of GSDMD is sufficient to mediate all steps necessary for membrane insertion, including the electrostatically driven membrane association, as well as the conformational changes that likely expose the extended beta sheet that dips into the plasma membrane. This for the first time allowed us to observe and study GSDMD membrane insertion in (living) human cells, supporting the conclusion that pores can grow monomer by monomer in a target membrane. These results are in line with previous *in vitro* findings in artificial membranes, showing that human GSDMD^NT^ assembles smaller arcs or slits, which grow into ring-shaped assemblies^4,15^. Likewise, atomistic molecular dynamics simulations predicted that small GSDMD^NT^ assemblies can already form ion-conducting membrane pores and provide a plausible pathway to pore opening in intact biolayers^48^. Earlier work had proposed the formation of GsdmA3 or GSDMD prepores composed of ring-like assemblies of GsdmA^NT^ or GSDMD^NT^ associated with membranes in the globular conformation, resembling the conformation of the N-terminal domain in full length gasdermin^12,13^. This model implied that a coordinated conformational change in all subunits gives rise to the eventual β-barrel structure that is inserted in the cell membrane^12,13^. Our data suggests that monomers of GSDMD^NT^ undergo conformational changes that allow membrane insertion, even if oligomerization is prohibited with nanobodies. This indicates that the assembly of prepores is not necessary for membrane insertion and therefore unlikely to be critical for pore formation. Recent atomic force microscopy data on GsdmA3 pores experimentally confirms membrane penetration of growing pores in different morphologies *in vitro*^16^. While mobile prepore-like assemblies were observed to attach to membranes and disappear, none of them were observed to penetrate the membrane. Nevertheless, findings in this and our work do not completely rule out that two different pathways to pore-formation exist in parallel^16^.

As the inhibitory nanobodies stabilize GSDMD into a monomeric intermediate, they can function as valuable tools to further elucidate molecular details of membrane insertion and pore formation in relevant cell types using live cell microscopy. In a parallel study, Kopp et al. solved the crystal structure of VHH_GSDMD-2_ and VHH_GSDMD-6_ in complex with full length GSDMD (Kopp et al., submitted)^49^. VHH_GSDMD-2_ binds to an epitope of full length GSDMD that forms the oligomerization interface in GSDMD pores, confirming the biochemically determined interference with oligomerization. VHH_GSDMD-2_ and VHH_GSDMD-1_ further competed for an overlapping epitope on GSDMD. Importantly, we found that fluorescent fusions of GSDMD^NT^ almost exclusively insert into the plasma membrane if oligomerization is inhibited by binding of antagonistic nanobodies. This demonstrates that the plasma membrane is indeed the primary target membrane of GSDMD^NT^ pores and mitochondrial localization of GSDMD^NT^ is unlikely during the first phase of pore formation in macrophages (Figure 3G). By extension, similar antagonistic nanobodies may reveal to which membranes the N-terminal effector domains of other gasdermin family members are targeted.

Internal structures containing fluorescent fusions of GSDMD^NT^ were observed in pyroptotic cells when pore formation was possible. Stabilization of monomers therefore allows us to distinguish the localization of GSDMD^NT^ before and after pore formation. We conclude that intracellular structures containing GSDMD^NT^ are a consequence of pore formation. They very likely constitute GSDMD pores removed from the plasma membrane by endocytic membrane repair processes, suggesting that membrane shedding is not the only mechanism to dispose GSDMD pores^35^, as previously described for MLKL^50^.

The antagonistic GSDMD nanobodies also provided new insights into the interconnectivity of the different cell death pathways in macrophages. Despite the presence of fully cleaved endogenous GSDMD in cells expressing VHH_GSDMD-1_ or VHH_GSDMD-2_, macrophages undergo apoptosis that is dependent on inflammasome assembly, ASC specks, and caspase-1 activity. Inflammasome-mediated apoptosis has previously been reported in caspase-1 knockout cells, in cells expressing catalytically inactive caspase-1, as well as in cells lacking GSDMD^1,41,51^. Caspase-1-mediated apoptosis in the absence of GSDMD pores is mechanistically different from the described ASC-dependent caspase-8-mediated apoptosis, which was observed in the explicit absence of caspase-1 activity^41,42^. Although ASC^PYD^ can nucleate polymerization of caspase-8 death effector domains (DEDs) in absence of caspase-1^40^, we found that caspase-3 activation was minimal in the absence of caspase-1, suggesting that direct recruitment of caspase-8 to ASC specks does not substantially contribute to the observed early apoptosis. Interestingly, apoptosis observed in our system followed a kinetic comparable to pyroptosis, with MxiH-stimulated cells already exhibiting caspase-3 activity as well as apoptotic morphology within 20-60 minutes after treatment. In contrast, canonical apoptosis, e.g., triggered by staurosporine, is a slower process in which caspase-3 activity is only emerging after three or more hours^52^.

Importantly, we report for the first time that inflammasome activation in the absence of GSDMD pore formation strongly augments caspase-1 activity. Only this enhanced activity resulted in efficient cleavage of caspase-3, caspase-7, and their substrates. We therefore propose a key regulatory role for the caspase-1 activity, which seems to be reduced when pores are formed. Formation of PFO pores of similar size also reduced the augmented caspase-1 activity, suggesting that caspase-1 activity is altered by ion fluxes or any other direct or indirect consequence of pore formation. This suggests that a so far elusive layer of regulation acts on caspase-1. It is possible that the most active form of cleaved caspase-1, the (p33/p10)_2_ form^53,54^, is stabilized in the absence of pores by preventing or delaying the secondary cleavage between the CARD and p20, which is associated with loss of activity. Pore formation may also provide some unidentified feedback signal to caspase-1 to dampen activity.

Remarkably, GSDME was efficiently cleaved in cells that assembled inflammasomes in the absence of GSDMD pores. Yet, we did not observe pyroptosis mediated by GSDME^NT^. This corroborates earlier findings that suggested that GSDME-induced lytic cell death does not play a major role in macrophages^55–57^. In contrast, overexpressed GSDME^NT^ in HEK293T cells^58,59^ as well as GSDME cleaved by caspase-3 in SH-SY5Y and MeWo cells^60^ are sufficient to initiate pyroptosis. This suggests that GSDME^NT^ may be subject to additional layers of regulation. Interestingly, GSDMD pore formation may also be further regulated: Efficient GSDMD pore formation relies on ROS generated by the Ragulator-Rag-mTORC1 pathway, which likely mediates the oxidative post-translational modification of GSDMD cysteine 191^10^. The same residue has recently been proposed to undergo palmitoylation to facilitate membrane association and full activation^61,62^.

Since the number of diseases in which the inflammasome and GSDMD play a detrimental role keeps increasing, there is a growing interest in specific GSDMD inhibitors^45^. Our proof-of-concept experiments therefore highlight the interesting therapeutic potential of antagonistic nanobodies VHH_GSDMD-1_ and VHH_GSDMD-2_. At the crossroads of intracellular signaling upon pathogenic threats and DAMPs, targeting GSDMD would not only prevent inflammation upon canonical but also non-canonical inflammasome stimuli. We could show that the extracellular addition of the nanobodies drastically reduces pyroptosis and the release of the pro-inflammatory cytokine IL-1β in both PMA-differentiated THP-1 macrophages as well as M-CSF-differentiated primary human macrophages. We propose that the nanobodies enter the cells upon the formation of the first sublytic GSDMD pores, rendering further GSDMD^NT^ oligomerization and thus pore formation and pyroptosis impossible (Figure 5F). One additional benefit of the therapeutic application of recombinant nanobodies is that the nanobodies in this scenario only target cells that have already assembled GSDMD^NT^ pores, i.e., nanobodies only gain access to cells relevant for the inflammatory response. The high specificity for both the target protein and the cellular state therefore renders antagonistic GSDMD nanobodies suitable for therapy. This is of particular interest as most other reported inhibitors of GSDMD are cysteine-reactive compounds and therefore may lack specificity^33,45,63,64^. The possibility to tailor nanobodies into bivalent or multivalent molecules may constitute a good basis for further optimizations^65^. Lastly, nanobody-mediated survival of macrophages with cleaved GSDMD likely relies on membrane repair processes, which could be studied in more detail using this system.

In conclusion, we show that antagonistic GSDMD nanobodies afford unprecedented modes of intervention by stabilizing informative intermediates of GSDMD^NT^ pore formation. The observed functional perturbation not only allowed mechanistic insights into membrane insertion and pore formation, but also provides an interesting proof of concept for the therapeutic application of recombinant nanobodies.

## Materials and methods

### Cell lines

Human embryonic kidney (HEK) 293T cells (ATCC CRL-3216, RRID: CVCL_0063), were cultivated in DMEM GlutaMax™ medium (Thermo Fisher Scientific) containing 10% FBS; THP-1 cells (ATCC TIB-202, RRID: CVCL_0006) were cultured in RPMI 1640 GlutaMax™ medium (Thermo Fisher Scientific) containing 10% FBS and 50 µM 2-mercaptoethanol. All genetically modified cell lines were generated by lentiviral transduction using lentivirus produced with packaging vectors psPax2 and pMD2.G (kind gifts from Didier Trono, École polytechnique fédérale de Lausanne, Switzerland). THP-1 or HEK293T cell lines constitutively expressing VHH_GSDMD-1_, VHH_GSDMD-2_, VHH_GSDMD-3_, VHH_NP-1_, or VHH_ASC_ with a C-terminal HA tag or C-terminal EGFP fusion under the control of the human elongation factor-1 α promoter (pEF1α) were generated using lentiviral vectors constructed by Gateway cloning (Thermo Fisher Scientific) using vectors modified from pRLL (a kind gift of Susan Lindquist, Whitehead Institute of Biomedical Research), followed by selection in 0.75 µg/mL puromycin (Thermo Fisher Scientific). Cell lines inducibly expressing the C1C-EGFP or C1C-mCherry inflammasome reporter were generated using lentiviruses produced with derivates of pInducer20 (a kind gift of Stephen Elledge, Harvard Medical School)^66^, followed by selection in 500 µg/mL geneticin (Thermo Fisher Scientific). These cell lines formed the basis for further lentiviral transduction to incorporate the constitutively expressing nanobodies as described above. THP-1 ΔASC cells were generated by lentiviral transduction with derivatives of pLenti CRISPR v2 (a kind gift from Feng Zhang, Broad Institute) with the targeting sequence GCTGGATGCTCTGTACGGGA. A representative single cell clone was validated by immunoblot and genomic DNA sequencing. Derivative THP-1 ΔASC cells expressing EGFP fusions of VHH_GSDMD-1_, VHH_GSDMD-2_, VHH_NP-1_ or VHH_ASC_ were generated by lentiviral transduction with derivatives of pRRL, followed by sorting for EGFP positive cells using a BD FacsAria Fusion cell sorter. All expression levels were verified by flow cytometry using the encoded fluorescent protein, or anti-HA staining with anti-HA B6 HA.11 (1:1000) and anti-mouse IgG Alexa Fluor 488 (1:500). Cells were fixed in 4% formaldehyde and measured using BD FACSCanto or Miltenyi MACSQuant flow cytometers. Cell lines are routinely tested for Mycoplasma contamination. All experiments involving lentiviruses were conducted in a Biosafety Level 2 laboratory.

### Primary cells

Human CD14^+^ monocytes were isolated from human whole blood buffy coats obtained from the blood bank of the University Hospital Bonn, with consent of healthy donors and according to protocols accepted by the institutional review board of the University of Bonn (#107/16). PBMCs were isolated using Ficoll-Paque™ PLUS (VWR) according to the manufacturer’s suggestions and monocytes purified using positive selection with paramagnetic CD14 (human) MicroBeads (Miltenyi Biotec). CD14^+^ monocytes were differentiated into macrophages using 100 ng/mL of recombinant human M-CSF (Immunotools) or 500 U/mL of recombinant human GM-CSF (Immunotools) in RPMI 1640 GlutaMax™ medium supplemented with 10% FBS, 500 U/mL PenStrep, and 1 mM sodium pyruvate for 3 days. To express VHH_GSDMD-1_, VHH_GSDMD-2_, or VHH_NP-1_ in combination with C1C-EGFP in primary macrophages, cells were transduced with the lentivirus packaging fusion protein Vpx-Vpr to counteract SAMHD1 restriction. To produce lentivirus, HEK293T cells were transfected with psPax2, pMD2.G, pCAGGS Vpx-Vpr, and lentiviral vectors based on pInducer20bi-NA, a derivative of pInducer20-NA with the bidirectional doxycycline-inducible promoter from pTRE3G-BI (TaKaRa). Expression of VHH-HA and the C1C-EGFP inflammasome reporter was thus doxycycline-inducible. Lentivirus was harvested 48 h post transfection, filtered through a 0.4 µm filter, and used to transduce primary macrophages in the presence of 10 µg/mL polybrene for 6 h. The next day, expression of both the VHH and the C1C-EGFP was induced with 1 µg/mL doxycycline for 24 h.

### Proteins

*Expression and purification of His-SUMO-GSDMD, His-SUMO, His-LFn-MxiH, and PA* Expression vectors for human His-SUMO-GSMD and His-SUMO were generated by inserting SUMO-GSDMD into pET28 by Gibson cloning. Proteins were expressed in *Escherichia (E.) coli* LOBSTR^67^ cells in Terrific Broth induced with 0.2 or 1 mM IPTG at an OD600 of 0.6. Cells were cultivated for 24 h at 18° C and lysed by French Press or sonication with a Bandelin Sonopuls HD2070 with TT13 tip. Subsequently, the proteins were purified by Ni-NTA affinity chromatography using Ni-NTA agarose beads (Qiagen) and gel filtration with a HiLoad 16/600 Superdex 75 pg column in buffers containing 20 mM HEPES pH 7.4, 150 mM NaCl, 10% glycerol, and 1 mM DTT. *B. anthracis* protective antigen (PA) and the fusion of *B. anthracis* LFn (aa 1-255) and *Shigella flexneri* MxiH were expressed in *E. coli* BL21(DE3) and purified as described before^25^.

### Expression and purification of nanobodies

Coding sequences for the different GSDMD nanobodies and the control VHH_NP-1_ were cloned into pHEN6-based bacterial, periplasmic expression vectors with C-terminal LPETG-His_6_ (large scale) or HA-His_6_ (small scale) tags using Gibson cloning. *E. coli* WK6 bacteria were transformed with nanobody expression vectors and grown in Terrific Broth. Expression was induced with 1 mM IPTG at an OD_600_ of 0.6, followed by cultivation at 30° C for 16 h. Bacterial pellets were resuspended in TES buffer (200 mM Tris-HCl pH 8.0, 0.65 mM EDTA, 0.5 M sucrose), after which periplasmic extracts were generated by osmotic shock in 0.25x TES at 4°C overnight. Nanobodies were purified with Ni-NTA agarose beads (Qiagen), followed by desalting with PD MiniTrap G-25 columns (GE Healthcare Life Sciences) (ELISA experiments) or gel filtration with a HiLoad 16/600 Superdex 75 pg column (tissue culture experiments) in buffers containing 20 mM HEPES pH 7.4, 150 mM NaCl, and 10% glycerol. For experiments with primary cells, endotoxin was removed using the Pierce™ High Capacity Endotoxin Removal Spin Columns (Thermo Fischer Scientific).

### Antibodies

The following antibodies were used: rabbit polyclonal anti-BID (Cell Signaling Technology Cat#2002S, RRID:AB_10692485), rabbit anti-caspase-3 clone D3R6Y (Cell Signaling Technology Cat#14220, RRID:AB_2798429), rabbit anti-cleaved caspase-3 (Asp175) clone 5A1E (Cell Signaling Technology Cat#9664S, RRID:AB_2070042), rabbit anti-caspase-7 clone D2Q3L (Cell Signaling Technology Cat#12827T, RRID:AB_2687912), mouse anti-caspase-8 clone 1C12 (Cell Signaling Technology Cat#9746S, RRID:AB_2275120), mouse anti-caspase-9 clone C9 (Cell Signaling Technology Cat#9508S, RRID:AB_2068620), rabbit anti-DFNA5/GSDME clone EPR19859 (Abcam Cat#ab215191, RRID:AB_2737000), rabbit polyclonal anti-E-tag-HRP (Bethyl Cat#A190-133P, RRID:AB_345222), mouse anti-GAPDH clone 0411 (Santa Cruz Biotechnology Cat#sc-47724, RRID:AB_627678), rabbit polyclonal anti-GSDMD (Atlas Antibodies Cat#HPA044487, RRID:AB_2678957), mouse anti-HA.11 Epitope tag clone 16B12 (BioLegend Cat# 901503, RRID:AB_2565005), mouse anti-HA-HRP clone 6E2 (Cell Signaling Technology Cat# 2999S, RRID:AB_1264166), goat polyclonal anti-mouse IgG (H+L)-HRP (Invitrogen Cat#31430, RRID:AB_228307), goat polyclonal anti-rabbit IgG (H+L)-HRP (Invitrogen Cat#31460, RRID:AB_228341), highly cross-adsorbed goat polyclonal anti-mouse IgG (H+L)-Alexa Fluor^TM^ 488 (Thermo Fisher Scientific Cat#A-11029, RRID:AB_2534088), highly cross-adsorbed goat polyclonal anti-rabbit IgG (H+L)-Alexa Fluor^TM^ Plus 647 (Thermo Fisher Scientific Cat#A32733, RRID:AB_2633282), rabbit anti-PARP clone 46D11 (Cell Signaling Technology Cat#9532S, RRID:AB_659884), mouse anti-TOM20 clone 29 (BD Biosciences Cat#612278, RRID:AB_399595), and mouse anti-vinculin clone hVIN-1 (Sigma-Aldrich Cat#V9131, RRID:AB_477629).

### Small compound inhibitors and reagents

The following small compound inhibitors and reagents were used: caspase-3/7 inhibitor I (Sigma), CRID3 (MCC-950) (Tocris), doxycycline (Biomol), LPS-EK Ultrapure (Invivogen), MG-132 (Selleckchem), Nigericin sodium salt (Biomol), PMA (phorbol 12-myristate 13-acetate) (Sigma Aldrich), Roche cOmplete™ Mini protease Inhibitor Cocktail (Sigma Aldrich), staurosporine (Enzo), Vx-765/belnacasan (Selleckchem), Z-VAD(Ome)-FMK (MedChemExpress).

### Nanobody library generation

To raise heavy chain-only antibodies (VHHs) against human GSDMD, a male alpaca was immunized four times with 200 µg GSDMD using Imject™ Alum Adjuvant (Thermo Fisher Scientific) according to protocols authorized by the MIT Institutional Animal Care and Use Committee. The VHH plasmid library in the M13 phagemid vector pD (pJSC) was generated as described before^25,68^. In brief, RNA from peripheral blood lymphocytes was extracted and used as a template to generate cDNA using three sets of primers (random hexamers, oligo(dT), and primers specific for the constant region of the alpaca heavy chain gene). VHH coding sequences were amplified by PCR using VHH-specific primers, cut with AscI and NotI, and ligated into an M13 phagemid vector (pJSC) linearized with the same restriction enzymes. *E. coli* TG1 cells (Agilent) were electroporated with the ligation reactions and the obtained ampicillin-resistant colonies were harvested, pooled, and stored as glycerol stocks.

### Nanobody identification by phage display

GSDMD-specific VHHs were obtained by phage display and panning with a protocol modified from Schmidt et al.^25^. *E. coli* TG1 cells containing the VHH library were infected with helper phage VCSM13 to produce phages displaying the encoded VHHs as pIII fusion proteins. Phages in the supernatant were purified and concentrated by precipitation. Phages presenting GSDMD-specific VHHs were enriched using chemically biotinylated GSDMD immobilized on Dynabeads™ MyOne™ Streptavidin T1 (Life Technologies). The retained phages were used to infect *E. coli* ER2738 and subjected to a second round of panning. 96 *E. coli* ER2837 colonies yielded in the second panning were grown in 96-well plates and VHH expression was induced with IPTG. VHHs leaked into the supernatant were tested for specificity using ELISA plates coated with control protein SUMO or SUMO-GSDMD. Bound VHHs were detected with HRP-coupled rabbit anti-E-Tag antibodies (1:10,000), and the chromogenic substrate tetramethylbenzidine (TMB) (Life Technologies). Reactions were stopped with 1 M HCl and absorption at 450 nm was recorded using a SpectraMax i3 instrument and the SoftMax Pro 6.3 Software (Molecular Devices). Positive candidates were sequenced and representative nanobodies were cloned into bacterial and mammalian expression vectors for further analysis.

### Nanobody ELISA

To test nanobody candidates, SUMO-GSDMD or SUMO in PBS were immobilized on ELISA plates at a concentration of 1 μg/mL overnight. Subsequently, the immobilized antigens were incubated with the HA-tagged nanobodies in 10% FBS/PBS in a 10-fold dilution series ranging from 100 nM to 1 pM. The nanobodies were detected using the mouse anti-HA HRP antibody (1:5000) and developed using the chromogenic substrate TMB. The reaction was stopped using 0.5 M HCl, after which the absorption was measured at 450 nm using a SpectraMax i3 instrument and the SoftMax Pro 6.3 Software (Molecular Devices).

### LUMIER assay

To test the functionality of VHHs in the reducing environment of the cellular cytosol, LUMIER assays were performed. 2.5·10^5^ HEK293T cells per well were seeded into 24-well plates, and were co-transfected the next day with 0.25 µg pCAGGS VHH-HA expression vectors and 0.25 µg of pcDNA3.1-based expression vectors for the Renilla-fused bait proteins GSDMD, GSDMD 4A, GSDMD^NT^ 4A, GSDMD^CT^ or the control NLRP1^CARD^ using PEI Max (Polysciences). High-binding Lumitrac 600 white 96-well plates (Greiner) were coated with 20 µg/mL of the mouse anti-HA.11 Epitope tag clone 16B12 antibody in PBS. One day post transfection, HEK293T cells were lysed in LUMIER lysis buffer (50 mM Hepes-KOH pH 7.9, 150 mM NaCl, 2 mM EDTA, 0.5% Triton X-100, 5% glycerol and Roche cOmplete™ Mini protease Inhibitor Cocktail) and bound to anti-HA-coated Lumitrac 600 plates for one hour to immunoprecipitate (IP) VHH-HA. After repeated washing, Renilla luciferase substrate coelenterazine-h was added to the IP well or lysate controls. Luminescence was measured using a SpectraMax i3 instrument and the SoftMax Pro 6.3 Software (Molecular Devices). The values plotted are the IP luminescence values normalized by the values of the lysate.

### Cell death quantification by LDH release

To quantify pyroptotic cell death, THP-1 cells were differentiated with 50 µg/mL PMA for 18 h, followed by a 24 h resting period; primary human macrophages were differentiated with M-CSF as described above. 3·10^5^ cells were seeded in 24-well plates and intracellular VHH expression was induced with 1 µg/mL dox where inducible promoters were used. The NLRP3 inflammasome was activated by consecutive treatment with 200 ng/mL ultrapure LPS for 3 h, and 10 µM nigericin (Nig), a potassium ionophore derived from *Streptomyces hygroscopicus*, in OptiMEM for 1 h. Where indicated, 40 µM VX or 2.5 µM CRID3 were added 30 minutes before nigericin stimulation as well as with the stimulus. The NLRC4 inflammasome was activated using 1.0 µg/mL recombinant *B. anthracis* PA and 0.1 µg/mL LFn-MxiH in OptiMEM for 1 h^69^, where indicated in presence of 40 µM VX. The extracellular administration of recombinant VHHs in increasing concentrations (2, 20, 50, 100, and 200 µg/mL) occurred simultaneously with the inflammasome stimulus. To measure pyroptotic cell death in HEK293T cells, 5·10^5^ cells per well were seeded into 24-well plates and co-transfected the next day with expression vectors for VHH-HA (0.5 µg) and GSDMD^NT^ (0.25 µg) or empty vector using Lipofectamine™ 2000 (L2000) (Thermo Fisher Scientific). Supernatants were collected 24 h after the transient transfection. Lactate dehydrogenase (LDH) in the supernatants from either cell type was quantified using the LDH Cytotoxicity Detection kit (TaKaRa or Roche) according to the manufacturer’s instructions. Absorption at 492 nm was measured using a SpectraMax i3 instrument and the SoftMax Pro 6.3 Software (Molecular Devices). Medium background signals were subtracted from all values. Control samples, in which cells were lysed in 1% Triton X-100, were subsequently used to normalize LDH release.

### Cytokine quantification by HTRF

To quantify IL-1β secretion, supernatants obtained as described for the LDH release assays were subjected to human IL-1β Homogeneous Time Resolved Fluorescence (HTRF) assays (Cisbio) according to the manufacturer’s instructions. Samples were excited at 340 nm and emissions at 616 nm and 665 nm were measured using a SpectraMax i3 instrument. IL-1β levels were calculated by the SoftMax Pro 6.3 Software (Molecular Devices) based on the standard curve.

### Cell death quantification by DRAQ7 uptake

To quantify cell death over time, 4·10^4^ THP-1 cells per well were seeded in 96-well plates in the presence of PMA and treated as described above for the LDH assay. Medium with stimuli was complemented with the non-cell permeable DNA dye DRAQ7 (100 nM) (Biolegend). DRAQ7 uptake was analyzed using the Incucyte Live-Cell Imaging system (Sartorius). The cells were recorded every 5 minutes for a total of 5 h using the Incucyte SX5 instrument, taking 4 images per well. The number of DRAQ7-positive nuclei (cell death count) and the cell confluency were analyzed using the Incucyte 2021C software. For every single image, the cell death count was corrected by subtraction of the value at the start of the experiment. The corrected cell death count was further normalized to the cell confluency and average values from all 4 images were calculated and plotted over time.

### Quantification of expression levels, inflammasome assembly, and caspase-3 cleavage by flow cytometry

Transduction of primary human M-CSF macrophages with lentiviruses encoding the C1C-EGFP inflammasome reporter and VHH-HA was assessed by quantifying the fraction of cells positive for C1C-EGFP by flow cytometry. To estimate the concentration of intact cells per sample, cells were measured by flow cytometry for a fixed time period of 30 s. The reduction of cells per volume served as an indirect indication for pyroptotic cell death. To quantify C1C-EGFP specks as a proxy for inflammasome assembly, we first gated cells positive for C1C-EGFP (Area), and then plotted height against width of the C1C-EGFP signal. Cells with C1C-EGFP specks appeared as a separated population with enhanced height and reduced width as described elsewhere^29,32^. For these experiments, 1·10^5^ primary macrophages in 24-wells were lentivirally transduced and treated as described above. The NLRC4 inflammasome was activated using 1.0 µg/mL recombinant *B. anthracis* PA and 0.1 µg/mL LFn-MxiH for 1 h. The cells were harvested by trypsinization, fixed in 4% formaldehyde, and analyzed using a BD FACSCanto flow cytometer.

For the quantification of inflammasome assembly in presence of the VHH-EGFP fusions, PMA-differentiated THP-1 macrophages expressing both VHH-EGFP and C1C-mCherry were stimulated with either 1.0 µg/mL recombinant *B. anthracis* PA and 0.1 µg/mL LFn-MxiH, or with 200 ng/mL LPS for 3 h and 10 µM nigericin for 1 h. To prevent the loss of responding cells by caspase-1-dependent pyroptosis, the cells were stimulated in presence of 40 µM VX. The fraction of specking C1C-mCherry positive cells was measured with a BD LSRFortessa SORP flow cytometer. Experiments were also performed in absence of VX and revealed that pyroptotic cells are lost during sample processing, while untreated and apoptotic cells could be analyzed by flow cytometry.

To measure cleaved caspase-3 in PMA-differentiated THP-1 macrophages, we performed experiments as described for apoptotic cell death. 3·10^5^ THP-1 macrophages in wells of 24-well plates were treated with 1.0 µg/mL recombinant *B. anthracis* PA and 0.1 µg/mL LFn-MxiH for 1 h in presence of 40 µM VX where indicated. Staurosporine is a non-selective inhibitor of several kinases and was added for 20 h as a positive control for intrinsic apoptosis and caspase-3 activation^38,70^. After fixation, cells were stained with rabbit anti-cleaved caspase-3 primary antibody (1:2000) and goat anti-rabbit Alexa Fluor Plus 647-coupled secondary antibody (1:500) in Intracellular Staining Permeabilization Wash Buffer (Biolegend). The fraction of cells positive for cleaved caspase-3 was measured with a BD LSRFortessa SORP flow cytometer. All flow cytometry data was analyzed using the FlowJo 10.7.1 software.

### Confocal microscopy

For live cell confocal microscopy experiments, 9·10^4^ PMA-differentiated THP-1 cells or 2·10^4^ GM-CSF differentiated primary human macrophages were cultured in 15 µ-slide 8 well Ibidi chambers or black, clear bottom, TC treated PhenoPlate™ 96-well microscopy plates (Perkin Elmer), respectively. The NLRC4 inflammasome was activated using 1.0 µg/mL recombinant *B. anthracis* PA and 0.1 µg/mL LFn-MxiH for 1 h in imaging medium (RPMI with 10% FBS, 50 µM 2-mercaptoethanol, 30 mM HEPES, no phenol red). HEK293T cells constitutively expressing VHH-EGFP fusions were seeded in Ibidi chambers (9·10^4^ cells per well) coated with poly-L-lysine (mol wt 70,000-150,000) (Sigma Aldrich). They were transiently transfected with 0.25 µg of the expression vectors GSDMD I104N-mCherry or GSDMD^NT^ I104N-mCherry. 5 h post transfection, images were recorded at least every 10 minutes using the HC PL APO CS2 63×1.20 water objective on a Leica SP8 Lightning confocal microscope (37° C, 5% CO_2_). Images were processed using ImageJ 2.3.0 software. To stain endogenous proteins in microscopy samples, cells were seeded and treated as above, fixed in 4% formaldehyde in PBS for 20 minutes, permeabilized with 0.5% Triton X-100, and stained with Hoechst 33342 (Thermo Fisher Scientific) as well as mouse anti-TOM20 antibody (1:500) and goat anti-mouse IgG AF647 (1:1,000) in PBS + 10% goat serum.

### Immunoblot

To detect the presence and/or cleavage of proteins of interest, PMA-differentiated THP-1 cells were treated with 1.0 µg/mL recombinant *B. anthracis* PA and 0.1 µg/mL LFn-MxiH to activate the NLRC4 inflammasome. 3-4·10^5^ cells (in 24-well plates) or 1.25·10^6^ cells (in 6-well plates) were lysed in 100 µL or 300 mL RIPA buffer (50 mM Tris pH 7.4, 150 mM NaCl, 1% IGEPAL CA-630, 0.25% Na-deoxycholate, 2 mM EDTA, 0.1% SDS, Roche cOmplete™ Mini protease Inhibitor Cocktail), respectively. Samples were freshly supplemented with 4x SDS-PAGE buffer (yielding 50 mM Tris pH 6.8, 0.01% bromophenol blue, 10% glycerol, 2% SDS final) with or without 100 mM DTT, sheared and heated to 95°C for 5 minutes. Proteins were separated by SDS-PAGE with 10% or 12% gels. Separated proteins were transferred to PVDF membranes (0.45 mm, Merck) by semi-dry transfer. All immunoblots were blocked in blocking buffer (5% non-fat dry milk (NFDM) in TBS with 0.05% Tween-20) for ≥ 2 h and probed with the following primary antibody dilutions: anti-BID (1:500), anti-caspase-3 (1:500), anti-caspase-7 (1:500), anti-caspase-8 (1:500), anti-caspase-9 (1:500), anti-GAPDH (1:1000), anti-GSDMD (1:500), anti-GSDME (1:500), anti-PARP (1:500), anti-vinculin (1:1000). Membranes were incubated with primary antibodies in blocking buffer at 4° C overnight, washed, and probed with HRP-coupled secondary antibodies in blocking buffer (1:3000) for 2 h. Chemiluminescent signal was induced by Western Lightning® Plus-ECL (Perkin Elmer), except for immunoblots of BID, caspase-3, caspase-7, caspase-8, caspase-9, and GSDMD which required Western Lightning Ultra (Perkin Elmer). The signal was detected using a Fusion Advancer imaging system (Vilber) and images were taken using the EvolutionCapt SL6 software (Vilber).

### Caspase-Glo activity and cell titer blue assay

To quantify the activity of caspase-1, caspase-3/7, and caspase-8, 5·10^4^ PMA-differentiated THP-1 cells cultured in 96-well plates were stimulated for 1 h with 1.0 µg/mL recombinant *B. anthracis* PA and 0.1 µg/mL LFn-MxiH in OptiMEM. Next, the cells plus supernatants were combined with an equal volume of the respective caspase-Glo reagent for 1 h according to the manufacturer’s instructions (Promega). The mixture was then transferred to a Lumitrac 600 plate and luminescence was measured using a SpectraMax i3 instrument and the SoftMax Pro 6.3 Software (Molecular Devices). Caspase-1-Glo assays were performed in the presence of 60 µM MG-132. The luminescence value of the negative control (OptiMEM plus caspase-Glo reagent control) was subtracted from all measured values. In parallel, cell titer blue (CTB) assays were conducted to determine the viability of the cultured, untreated cells using the CellTiter-Blue® Reagent (Promega) according to the manufacturer’s instructions. Here, supernatants were aspirated and 100 µL of the CTB reagent was added to the cells in the 96-well plate for 1 h at 37°C. Samples were excited with light at a wavelength of 560 nm and fluorescence measured at 585 nm. The value for the control well without cells was subtracted and the cell line expressing the control VHH_NP-1_ was set to 1. The final CTB values were used to normalize for differences in cell numbers between the different cell lines in the caspase-Glo assays. For the caspase-1 Glo assay, values were further normalized to MxiH-treated VHH_NP-1_ samples.

## Acknowledgments

We would like to acknowledge the support of the Flow Cytometry Core Facility of the Medical Faculty, University of Bonn, for their support, services and devices funded by the Deutsche Forschungsgemeinschaft (DFG, German Research Foundation) for project number 16372401, 387335189, 387333827, and 216372545. We are grateful to Gabor Horvath and the Microscopy Core Facility of the Medical Faculty at the University of Bonn for providing help, services and instrumentation supported by the Deutsche Forschungsgemeinschaft (DFG, German Research Foundation) for project number 388159768, and the Bundesministerium für Bildung und Forschung (BMBF, Federal Ministry of Education and Research) (ACCENT; project number 01EO2107. We would like to thank Christina Martone and Jessica Ingram for providing the infrastructure for nanobody generation at the Whitehead Institute.

## Funding agencies

The presented work was supported by the following funding agencies:

Deutsche Forschungsgemeinschaft (DFG, German Research Foundation) grant SFB1403-414786233 (FIS)

Deutsche Forschungsgemeinschaft (DFG, German Research Foundation) Emmy Noether Programme 322568668 (FIS)

Deutsche Forschungsgemeinschaft (DFG, German Research Foundation) Germany’s Excellence Strategy – EXC2151–390873048 (FIS and MG)

## Author contributions

**Lisa D.J. Schiffelers:** Conceptualization, Data curation, Methodology, Investigation, Resources, Writing – original draft / review and editing, Visualization, Supervision; **Sabine Normann:** Methodology, Investigation; **Sophie C. Binder:** Investigation; **Elena Hagelauer:** Investigation; **Anja Kopp:** Conceptualization, Resources; **Assaf Alon:** Resources; **Matthias Geyer:** Funding acquisition, Resources; **Hidde L. Ploegh:** Funding acquisition, Resources; **Florian I. Schmidt:** Conceptualization, Data curation, Funding acquisition, Investigation, Methodology, Supervision, Writing – original draft / review and editing.

---

**Supplementary Fig. 1.**
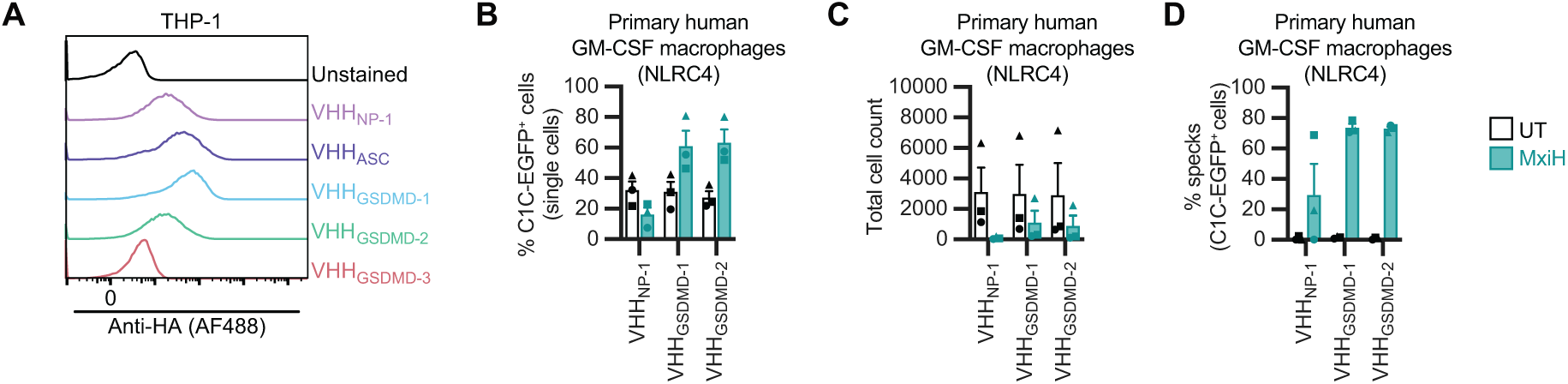
VHH_GSDMD-1_ and VHH_GSDMD-2_ abrogate pyroptosis. **(A)** THP-1 cell lines constitutively expressing the indicated HA-tagged nanobodies were fixed, stained for HA, and histograms of the HA signals of a representative experiment were displayed. **(B-D)** Primary GM-CSF-differentiated monocyte-derived human macrophage were transduced and stimulated as described in Fig. 2H-K. 1 h post treatment, cells were harvested, fixed, and analyzed by flow cytometry to determine the fraction of C1C-EGFP^+^ and thus VHH-expressing cells (B), cell count over 30 s (C), and the fraction of C1C-EGFP^+^ cells assembling ASC specks (D). Data represent average values (with individual data points) from three independent donors ± SEM.

**Supplementary Fig. 2.**
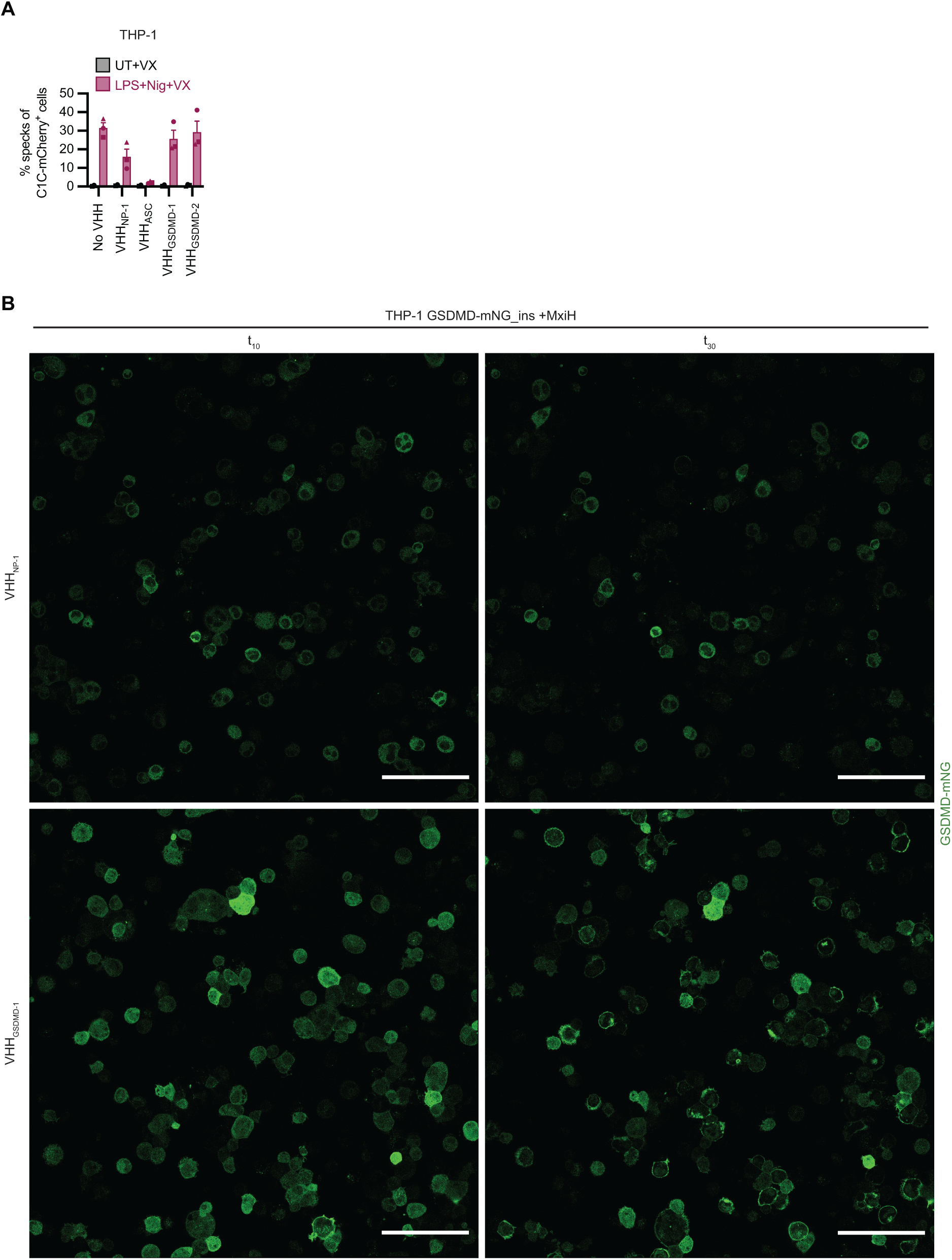
VHH_GSDMD-1_ and VHH_GSDMD-2_ do not interfere with inflammasome assembly. **(A)** THP-1 cell lines expressing C1C-mCherry (dox-inducible) as well as the indicated VHH-EGFP fusions (constitutively) were differentiated with PMA, treated with dox for 24 h, and stimulated with 200 ng/mL ultrapure LPS for 3 h and 10 µM nigericin (Nig) in presence of VX for 1 h to activate NLRP3. Cells were harvested and ASC specks were quantified by flow cytometry. Data represent average values (with individual data points) from three independent experiments ± SEM. **(B)** PMA-differentiated THP-1 cells constitutively expressing GSDMD-mNeonGreen_ins and the indicated HA-tagged nanobodies were stimulated with MxiH for 1 h as described in Fig. 2B. Representative images 10 or 30 minutes post treatment (t_10_ and t_30_) with larger fields of view from the same experiment shown in Figure 2B are displayed. Scale bar = 100 µm.

**Supplementary Fig. 3.**
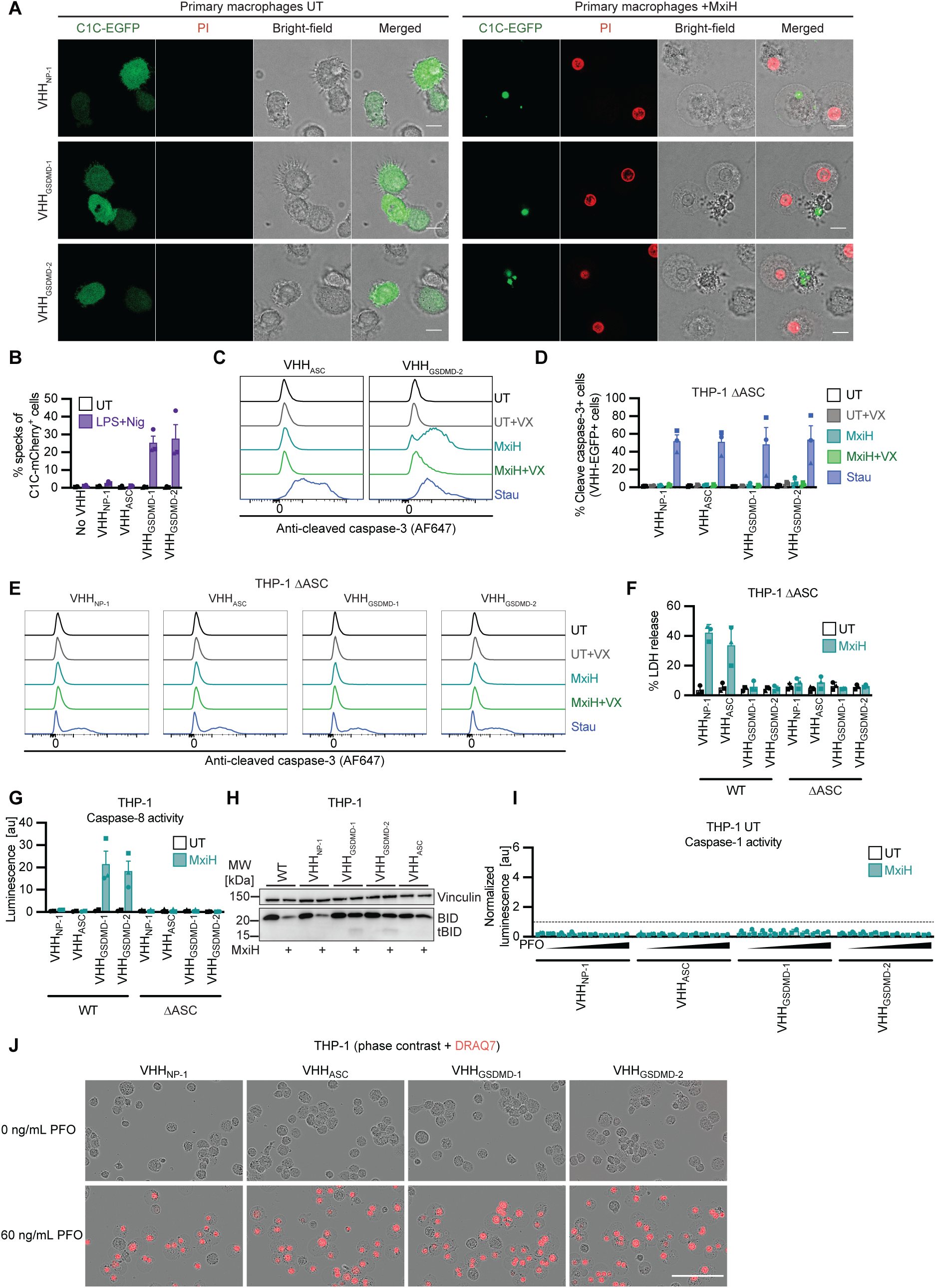
Inhibition of pore formation by antagonistic GSDMD nanobodies augments caspase-1 activity and triggers caspase-1-dependent apoptosis. **(A)** Primary GM-CSF-differentiated monocyte-derived human macrophages were transduced and stimulated as described in 2H-K. Cells were recorded by live-cell confocal microscopy and images representative of three independent donors are displayed. Scale bar = 10 µm. **(B-C)** PMA-differentiated THP-1 cell lines expressing C1C-mCherry (dox-inducible) as well as the indicated VHH-EGFP fusions (constitutively) were differentiated with PMA and treated with dox for 24 h. (B) Cells were stimulated with LPS for 3 h and nigericin (Nig) for 1 h to activate NLRP3 as described in Fig. 2C, harvested, and analyzed by flow cytometry to quantify C1C-mCherry specks as described in Fig. 4B. (C) Cells were stimulated with LFn-MxiH and PA (MxiH) for 1 h to activate NLRC4 as described in Figure 2B, stained for cleaved caspase-3, and analyzed by flow cytometry. Average data of three independent experiments are displayed in Fig. 4C; representative histograms of individual cell lines from the same experiment are shown in Fig. 4D and here. **(D-E)** PMA-differentiated THP-1 ΔASC cells constitutively expressing the indicated HA-tagged nanobodies were stimulated with MxiH for 1 h as described in Fig. 2B. 5 µM staurosporine (Stau) for 20 h was used as positive control to trigger apoptosis. Stimulation was performed in the absence or presence of VX as indicated. Cells were harvested, stained for cleaved caspase-3, and the fraction of cells positive for cleaved caspase-3 quantified by flow cytometry (D). Representative histograms of the staining for cleaved caspase-3 for indicated treatments are presented in (E). **(F-J)** PMA-differentiated THP^-^1 WT (G-J) or ΔASC (F,G) cells constitutively expressing the indicated HA-tagged nanobodies were stimulated with MxiH for 1 h as described in Fig. 2B (F-H) or treated with increasing PFO concentrations (0, 5, 10, 20, 30, 60, 120, 240, and 480 ng/mL) (I,J). LDH release was measured and normalized to cells lysed in Triton X-100 (F). Cells and supernatants were harvested to measure caspase-8 (G) or caspase-1 (I) activity using Caspase-Glo assays and activity was corrected for cell numbers per sample using CTB values (G,I). Caspase-1 activity was normalized to MxiH-treated cells expressing VHH_NP-1_ (indicated as dashed line) (I). Values for PFO-treated samples in the absence of MxiH are displayed in (I), while the corresponding data of MxiH-stimulated cells from the same experiments are displayed in Fig. 4L. Cell lysates were separated by SDS-PAGE and analyzed by immunoblot with the indicated antibodies (H); immunoblots representative of three independent experiments are displayed. THP-1 cells treated in the presence of DRAQ7 were recorded with an Incucyte Live-Cell Imaging system. Representative image of cells left untreated (0 ng/mL PFO) or treated with 60 ng/mL PFO after 30 minutes are displayed (J). Data represent average values (with individual data points) from three independent experiments ± SEM, unless otherwise indicated.

